# No single PCR test is sufficient to determine parvovirus IHHNV presence in or impact on farmed shrimp production

**DOI:** 10.1101/2023.09.08.556791

**Authors:** Kallaya Sritunyalucksana, Piyachat Sanguanrut, Jiraporn Srisala, Jumroensri Thawonsuwan, Nattakan Saleetid, Rapeepun Vanichviriyakit, Charoonroj Chotwiwatthanakun, Timothy W. Flegel, Suparat Taengchaiyaphum

**Author notes:** These authors with equal contribution to the work (Kallaya Sritunyalucksana, Piyachat Sanguanrut).

## Abstract

The main purpose of this report is to provide hard evidence that the shrimp parvovirus IHHNV has not resulted “in significant consequences e.g. production losses, morbidity or mortality at a zone or country level” in Thailand since at least 2010. It also reveals that no single PCR test is sufficient to identify IHHNV-infected shrimp. It presents historical evidence and new evidence from 11 commercial ponds cultivating the giant tiger shrimp *Penaeus monodon* in Thailand. These ponds were selected because they were the ponds that gave positive PCR test results for IHHNV using 2 methods recommended for IHHNV diagnosis by WOAH. However, an additional in-house “IHHNV Long-Amp method” (IHHNV-LA) was also used to amplify 90% of the 4 kb IHHNV genome sequence, and it also gave false-positive test results with 2 of the 11 ponds. Further tests using normal histopathological analysis for the presence of pathognomonic Cowdry A type inclusions (CAI), *in situ* hybridization (ISH) and immunohistochemistry (IHC) could confirm IHHNV infections in only 2 of the 4 ponds PCR-positive using all 3 PCR methods. In addition, positive detection of CAI alone was equivalent to ISH or IHC in confirming IHHNV infection after a positive test with any of the PCR methods used. In summary, the recommended WOAH PCR methods gave false positive test results for IHHNV infection with 9/11 ponds (82%). All 11 ponds gave profitable harvests despite the confirmation of IHHNV in 2 ponds, where it was accompanied by various additional pathogens. Unfortunately, according to current practice, positive PCR test results with the WOAH methods alone sometimes leads to rejection of traded shrimp products without assurance that the test results are not false positive results that may arise from endogenous viral elements (EVE).

## 1. INTRODUCTION

### 1.1. Identity of IHHNV

The official scientific name of infectious hypodermal and hematopoietic necrosis virus (IHHNV) is now Decapod *Penstylhamaparvovirus* 1 in the family *Parvoviridae* and sub-family *Hamaparvovirinae* (Pénzes et al., 2020). Acute, catastrophic disease with mass mortality was originally reported in IHHNV-infected *Penaeus stylirostris* (Lightner et al., 1983). It was discovered that the virus originated from grossly normal stocks of captured wild *Penaeus monodon* that were imported for experimental culture in the Americas in facilities where *P. stylirostris* was also under experimental culture. Subsequent IHHNV disease outbreaks in *Penaeus vannamei* in the Americas resulted not in mortality, but instead retarded growth accompanied by anatomical deformities (e.g., bent rostra) called runt deformity syndrome (RDS) (Bell and Lightner, 1984). The coefficient of variation (CV) in size for cultivated populations with RDS was typically 35-90%, with the mean shifted towards unmarketable small sizes that dominated crops and made them unprofitable. This compared to normal IHHNV-free populations with CVs of 10-30% (Bray et al., 1994, Brock and Main, 1994, Carr et al., 1996, Lightner, 1996, Primavera and Quinitio, 2000, Pruder G.D. et al., 1995). The effect of IHHNV on growth of *P. monodon* is controversial, but confirmed studies (Flegel et al., 2004, Withyachumnarnkul et al., 2006) indicate that it does not cause RDS or significant negative effects on growth similar to those that occur in *P. vannamei*.

IHHNV was among the 5 shrimp viruses listed in the first edition of the WOAH Aquatic Animal Health Code (AAHC) in 1995 and it is still listed in the current edition (WOAH, 2019b). Three distinct genotypes of IHHNV were listed in the 2019 online edition of the OIE Aquatic Manual of Diagnostic Tests for Aquatic Animals (WOAH, 2019a). These are Types 1 and 2 that are infectious viruses, while Type 3 was called non-infectious IHHNV. All are detectable by conventional polymerase chain reaction (PCR) tests. The primer pair 309F/R can detect all the known genotypes of infectious IHHNV while primer pair 389F/R can also detect non-infectious Type 3 (Tang et al., 2000, Tang et al., 2007) which comprises IHHNV-related DNA sequences present in the genome of *P. monodon* (Saksmerprome et al., 2011, Tang and Lightner, 2006, Tang et al., 2007). Type 3 has recently been revealed to be an assembly artifact that arose from scrambled sequences that sometimes occur in disjunctive clusters within the genome of *P. monodon* and constitute remnants of an ancient form of IHHNV (Huerlimann et al., 2022, Taengchaiyaphum et al., 2022). Such non-retroviral virus sequences in animal genomes are called endogenous viral elements (EVE) (Feschotte and Gilbert, 2012).

The target tissues for IHHNV infection in penaeid shrimp are of ectodermal origin (e.g., the gills and sub-cuticular epidermis), and mesodermal origin (e.g., haematopoietic tissue and antennal glands). Histological characteristic IHHNV lesions obtained using hematoxylin and eosin (H&E) staining are called Cowdry Type A inclusions (CAI). They comprise a central, eosinophilic, intranuclear inclusion separated by a small unstained space from marginated (basophilic) chromatin and are pathognomonic for IHHNV infection (Bell and Lightner, 1984, Lightner, 1996). However, CAIs are an artifact of the Davidson’s acid fixative that is recommended to preserve shrimp tissue for histological analysis, and they do not occur when other fixatives such as neutral phosphate buffered fixative are used. CAI are also produced during the early stages of infections with white spot syndrome virus (WSSV) but, unlike the inclusions of IHHNV, they cause massive nuclear enlargement, and they change to basophilic as they mature (Lightner, 1996).

### 1.2. Decline in negative economic impact of IHHNV

The negative impact of IHHNV on cultured shrimp production in the USA dropped dramatically after the development and adoption of domesticated, specific pathogen free (SPF) shrimp stocks free of the virus (Wyban et al., 1992, Wyban, 2007). This work in Hawaii revealed that stocking postlarvae (PL) derived from such shrimp in cultivation ponds did not result in significant RDS or other negative impact even if the shrimp became infected after stocking by residual virus from the previous culture cycle. Consistent, subsequent stocking of the same pond with IHHNV-free PL resulted in no negative impact and in a progressive decline in the number of IHHNV-positive specimens. Similarly, upon the first introduction of non-SPF *P. vannamei* for cultivation in Thailand in the late 1990’s a few cases of RDS were seen, but after the mid 2000’s in Thailand upon the widespread use of IHHNV-free PL of *P. vannamei*, no cases of RDS have been reported (Flegel, 2006).

In September 2020, a report from the OIE Aquatic Animal Health Standards Commission (AAHC) responded to an official request from Ecuador to review the impact of IHHNV and to consider removing it from its list of crustacean diseases [Report of the meeting of the OIE Aquatic Animal Health Standards Commission (AAHC), Paris, 19‒26 February 2020 in Section 4.6, Annex 9]. The AAHC upheld the continued listing of IHHNV solely because it concluded that IHHNV fit Criterion 4b: “The disease has been shown to affect the health of cultured aquatic animals at the level of a country or a zone resulting in significant consequences e.g. production losses, morbidity or mortality at a zone or country level”. If the AAHC had deemed that IHHNV did not satisfy this criterion, it would have been required to delist it.

To support their decision that IHHNV did satisfy Criterion 4b, the AAHC report cited 22 references, 8 of which were published prior to 2010 while only 14 covered the period 2010 to 2020 during which the impact of IHHNV on shrimp production should have been assessed. In **Supplementary Information 1**, these remaining 14 cited references are examined in detail, and it is shown that only 2 of them (Jagadeesan et al., 2019, Sellars et al., 2019) contained new data showing production losses in ponds infected with IHHNV, but they do not demonstrate mass stock loss or significant consequences as required by Citerion 4b. All the other 12 publications from 2010 onward concerned studies of IHHNV itself in terms of biology and detection and none contained original data regarding the impact of IHHNV on production.

The reasons why the remaining 2 publications are inappropriate to use to justify Criterion 4b are described in Supplementary Information 1. Both publications were based predominantly on positive PCR results for detection of IHHNV with no additional histological examination (e.g., *in situ* hybridization or immunohistochemistry) to confirm IHHNV infections. Whilst the data of Sellars et al. 2019 was taken from a broader data set where additional PCR was carried out to confirm samples were negative from all other endemic pathogens in the region, it is unknown if those of Jagadeesan et al. 2019 had any additional PCR data to demonstrate the presence of other pathogenic agents. Sellars et al. 2019 also based the financial cost calculations on an Australian production model which is unique compared to the rest of the world in that they achieve very high 8 tons/ha averages whilst the average globally is around 6.0-7.5 tons/ha. Clearly, all the harvests would have been profitable, but some more-so than others. Profits from farm to farm vary based on multiple, complex factors including management. Thus, it is not rational to refer to the profits of some farmers as losses because they are lower than those of another. In addition, for the paper by Jagadeesan et al. (2019), it was a survey of 350 *P*. *vannamei* farms in India of which 30 were PCR positive for IHHNV. Of these 16/30 had problems while 14/30 did not, indicating that IHHNV was not significantly associated with problem ponds and that other causes should have been sought. In addition, publications from the interval 2010 to 2020 that describe lack of significant problems linked to IHHNV were not included in the AAHC review (e.g., Hernández-Ruíz et al., 2020; Mendoza-Cano et al., 2014). Indeed, a recent publication by Aranguren Caro et al. (2022) showed a lack of significant impact of IHHNV on production is also documented, with IHHNV only causing small measurable reductions in growth in some ponds.

### 1.3. Study of arbitrarily selected Thai cultivation ponds positive for IHHNV by PCR

In this paper we present results from 11 recently harvested Thai shrimp ponds that were selected because they gave positive PCR test results for IHHNV using two WOAH PCR detection methods via active surveillance. The results also revealed significant problems might arise from false positive tests for IHHNV using currently recommended WOAH methods, if additional confirmatory tests are not carried out. In our experience, these should not be additional PCR tests, even with sequencing, but should be histopathological analysis for Cowdry A type inclusions (CAI) and/or *in situ* hybridization (ISH) or immunohistochemistry (IHC). We hope that this information will stir others to carry out similar studies and provide more evidence from production ponds to support the contention that IHHNV no longer has “significant consequences e.g. production losses, morbidity or mortality at a zone or country level.”

### 1.4 A precautionary note

It is important to understand from the outset that we do not consider IHHNV to be innocuous. It should not be ignored and must remain included in the group of pathogens that should be excluded from SPF shrimp used to supply shrimp farmers with PL to stock their cultivation ponds. The situation is the same as that for the viruses MBV and BP that were formerly listed by OIE but were later removed from the list (WOAH, 2008). They were removed because full understanding of their biology allowed for their exclusion from hatcheries and from post-larvae supplied to shrimp farmers for stocking in their cultivation ponds. Hepatopancreatic parvovirus (HPV), now called decapod hepanhamaparvovirus (DHPV), causes much more severe growth retardation than IHHNV (Flegel et al., 2004) but has never been listed for similar reasons. However, producers that use wild, captured PL or broodstock to produce PL must be aware that these unlisted pathogens must be excluded by screening broodstock or PL for their absence. This is because their presence may decrease production efficiency and profitability even though they may not cause financial losses.

The same situation now applies to IHHNV. The current, high-quality PL produced from SPF stocks of *P. vannamei* (Alday-Sanz et al. 2020) are not infected with IHHNV or other known pathogens and they produce offspring that can achieve average daily weight gains of 0.3 to 0.4 g or more under ideal culture conditions. In addition, they are unaffected by IHHNV infection. The weight gain is well beyond that possible with offspring obtained from captured, wild broodstock. This achievement has resulted from a combination of innovations in breeding, selection, feed and culture management. Thus, no pathogen can simply be ignored and assumed to have no possible negative impact on production. At the same time, no manageable shrimp pathogen should be considered as a perpetual dire threat when the evidence clearly shows that it is not.

Rational aquaculture management requires that shrimp farmers take precautions to ensure optimum culture conditions such as exclusion of known pathogens, including IHHNV. This can be achieved either by testing before stocking or by using PL from a supplier with a certified long history of providing PL free of known pathogens. For comparison, we pointed out from WHO data accessed on 10 December 2024 (https://data.who.int/dashboards/covid19/cases?n=c) that 180,907 world cases of Covid 19 were reported over 28 days up to 24 November 2024 together with 2,665 deaths. Thus, thousands more survive after suffering light to severe illness. All these Covid-19 infections, lethal or not, have negative impacts. Despite this, all restrictions on movements of people worldwide have now been removed. This does not mean that Covid-19 presents no threat so long as reasonable precautions are taken. In our opinion, the situation is similar to that for the shrimp viruses MBV, BP and HPV and the work described herein supports the argument that like them, IHHNV should not be a listed pathogen.

## 2. MATERIALS AND METHODS

### 2.1. Shrimp specimens

Juvenile giant tiger shrimp (*P. monodon*) were obtained from 2 sets of shrimp cultivation ponds. These ponds were selected based on being positive for IHHNV during an active surveillance using WOAH-recommended PCR methods. The PL used to stock the ponds were not tested for IHHNV, so the IHHNV source was unknown. The first, **Pond Set 1** consisted of 4 ponds, 1 from Chachoengsao province on the eastern coast of the Gulf of Thailand and 3 from Krabi province on the southwestern Andaman Sea coast of Thailand. These were cultivated from February to August, during the optimum time for shrimp production in Thailand. Another set of 7 ponds, **Pond Set 2** were obtained from the adjacent provinces Phuket and Phang Nga on the southwestern coast of Thailand on the Andaman Sea. These were cultivated from October to April, through the Thai winter (November to February) which is the riskiest time for shrimp cultivation in Thailand. However, this period attracts some farmers due to higher product prices.

Thirty shrimp were collected randomly from each pond at harvest to assess the range of shrimp sizes by fresh weight. For Pond Set 1, an additional 5 were collected from each pond for PCR and histological analysis. Pleopods from the 5 shrimp specimens were removed for PCR analysis, while the cephalothoraxes of the same specimens were processed using standard methods (Bell and Lightner, 1988) to confirm IHHNV presence by normal histological analysis plus ISH and IHC analysis. For Pond Set 2, the technician mistakenly prepared 6 shrimp specimens from each pond for PCR analysis and a separate set of 3 shrimp each for histological analysis. Thus, for Pond Set 2, results from the PCR tests and histological analysis could not be directly compared, but production results could be compared between ponds confirmed or not for the presence of infectious IHHNV.

### 2.2. DNA extraction

Pleopods were collected from 5 individual shrimp from each pond and homogenized for separate analysis in DNA lysis buffer (50 mM Tris-base, 100 mM EDTA, 50 mM NaCl, 2% (w/v) SDS and 100 µg/ml proteinase K). The mixtures were incubated at 56 °C for 1 h before DNA was extracted using an QIAamp DNA Mini Kit (Qiagen, Germany) according to the manufacturer’s protocol. Total DNA concentration was determined by Qubit 3.0 Fluorometer (Life Technologies, USA).

### 2.3. PCR detection methods for IHHNV

Three PCR detection methods for IHHNV detection were used in this study (**Table 1**). Two comprised WOAH-recommended methods here called the IHHNV-309 method employing primer pairs 309F/R and the IHHNV-389 method employing primer pairs 389F/R (Tang et al., 2000, Tang et al., 2007). These methods were carried out following the WOAH manual (WOAH, 2019a). A third, long-amp IHHNV (IHHNV-LA) detection method consisted of a two-step nested PCR protocol that was established in our laboratory to cover 90% (∼3.6 kb) of the whole genome sequence of infectious IHHNV (4.1 kb). The primers were designed from the GenBank database of IHHNV accession number AF218266. The sensitivity limit of the IHHNV-LA method (10 copies/ng total DNA) was determined using a DNA plasmid containing an insert of the 3,665 bp-product from the first amplification step as the template (Taengchaiyaphum et al., 2021).

**TABLE 1.**
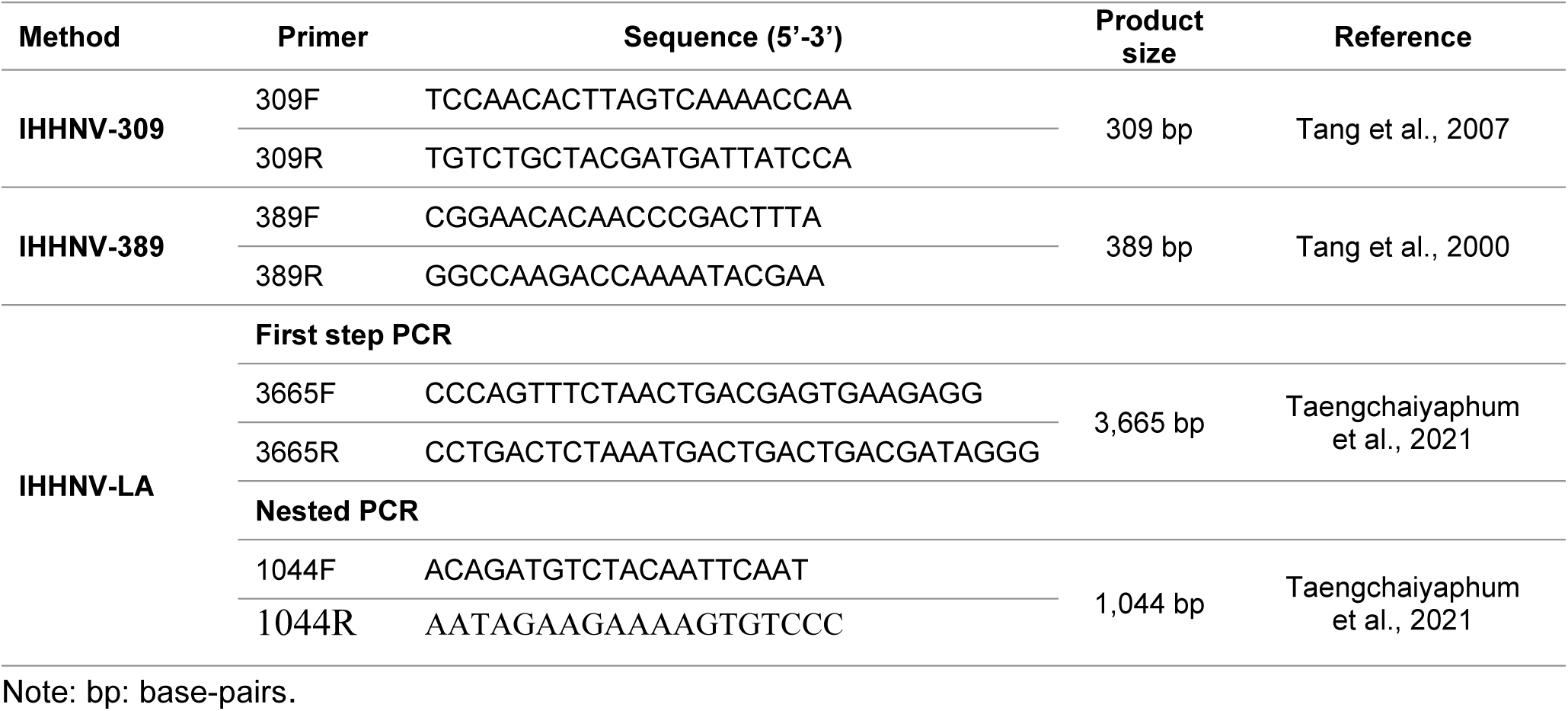
Primers used for detection of IHHNV by PCR in this study.

### 2.4. Histological analysis

Histological analysis was carried out by standard methods (Bell and Lightner, 1988). Briefly, the cephalothorax of each shrimp sample was fixed with Davidson’s fixative for 24 h and then changed to 70% ethanol. The tissue was next dehydrated with a series of graded ethanol ending with xylene and embedded in paraffin. Sections were cut at 4 µm and placed onto glass slides for H&E staining to detect of Cowdry Type A inclusions (CAI) characteristic for IHHNV, for *in situ* hybridization (ISH) analysis and for immunohistochemistry (IHC) analysis (Lightner, 1996). Positive control slides were prepared using archived tissue blocks of shrimp infected with IHHNV and showing typical IHHNV-CAI. We consider the detection of CAI as the gold standard for confirmation of IHHNV infection. The farm samples were also examined histologically for the presence of lesions in the hepatopancreas (HP) caused by monodon baculovirus (MBV), hepatopancreatic parvovirus (HPV) and other pathogens such as bacteria known to cause retarded growth in *P. monodon* (Flegel et al., 2004, Flegel, 2006). The viruses MBV and HPV are now called *Penaeus monodon* nudivirus (PmNV) (Yang et al., 2014) and *Penaeus monodon* hepandhamaparvovirus (Cotmore et al., 2019), respectively. Analysis of the lesions for these 2 viruses was carried out as recommended by Lightner (1996). The HP of the specimens were also screened for the presence of the microsporidian *Enterocytozoon hepatopenaei* (EHP) that is known to infect the HP and to be associated with retarded shrimp growth (Chaijarasphong et al., 2021).

### 2.5. *In situ* hybridization (ISH)

The PCR products amplified from both IHHNV-309 and IHHNV-389 methods were used as templates to prepare digoxigenin (DIG)-labeled IHHNV probes using a PCR DIG-labeling kit (Roche, Germany). After preparation, probes were purified using a Gel/PCR cleanup kit (Geneaid, Taiwan) using the protocol described by the manufacturer. The protocol for ISH has been previously described (Sanguanrut et al., 2022). Briefly, slides containing adjacent tissue sections to those used for histological analysis by H&E staining were de-paraffinized, rehydrated and TNE buffer (500 mM Tris-Cl, 100 mM NaCl, l0 mM EDTA) before treatment with proteinase K (Sigma, USA) and assay using 200 ng per slide of the DIG-labeled probes in hybridization buffer. A control reaction without probe was included in a separate container. After incubation overnight at 42 °C the slides were washed, incubated with blocking buffer and further incubated with 1:500 anti-DIG-AP antibody in the dark, washed and developed for signal using NBT/BCIP solution (Roche, Germany) and counterstained with 0.5% Bismarck Brown Y (Sigma, USA) for microscopic examination (Leica DM750, ICC50W model with a digital camera and LASV4.12 Software).

### 2.6. Immunohistochemistry (IHC) analysis

A polyclonal antibody to the capsid protein of IHHNV was produced in a rabbit using recombinant IHHNV capsid protein and the concentration appropriate for IHC was determined. The tissue sections were de-paraffinized, rehydrated with xylene and a series of graded alcohols to water. They were then treated with 0.6% H_2_O_2_ in methanol for 15 min to quench endogenous peroxidase activity followed by washing two times with 1X PBS for 5 min each. The slides were incubated with blocking buffer (10% fetal bovine serum in 1X PBS pH 7.4) for 1 h, at RT and replaced with rabbit anti-capsid IHHNV antibody (1:1000 in blocking buffer) (Kiatmetha et al., 2018) in a moist chamber at 37 °C for 1-2 h. The slides were washed three times to remove unbound antibody with 1X PBS for 5 min each followed by incubation with horseradish peroxidase (HRP)-conjugated goat anti rabbit antibody (1:500 in the blocking buffer) at 37 °C for 1 h in a moist chamber before washing three times with 1X PBS for 5 min each. The slides were incubated with a VECTOR^®^NovaRED^TM^ substrate kit (Vector laboratories, USA) for 5-15 min and washed with water for 5 min before counterstaining with Mayer’s hematoxylin, dehydrating in a series of graded alcohols to xylene and covering with a cover glass.

## 3. RESULTS

### 3.1. History of official surveillance reports of IHHNV in Thailand

For general interest, the percentage prevalence of IHHNV in samples of both *P. monodon* and *P. vannamei* received by the Thai Department of Fisheries (Thai DOF), the competent authority for shrimp diseases in Thailand from the years 2011 to 2021 is shown in **Figure 1** together with National production. In 2013, production dropped precipitously due mostly to reduced cultivation because farmers feared acute hepatopancreatic necrosis disease (AHPND) (Sanguanrut et al., 2018). Subsequent recovery to pre-2013 levels has not occurred due to bankruptcy of the majority of Thai shrimp export companies during the AHPND crisis, and to concomitant loss of export market share to other shrimp producing countries. In any case, the graph clearly shows that the percentage of IHHNV positive samples was not significantly correlated with a negative effect on shrimp production. In fact, a Pearson correlation analysis (**Figure 2**) showed a significant (p = 0.015) positive association between production quantity and the percentage of positive IHHNV detections in the total number of farmed shrimp specimens examined by active and passive surveillance by the Thai DOF. This result clearly reveals that simple data on detection of IHHNV prevalence cannot be used to predict shrimp production level.

**FIGURE 1.**
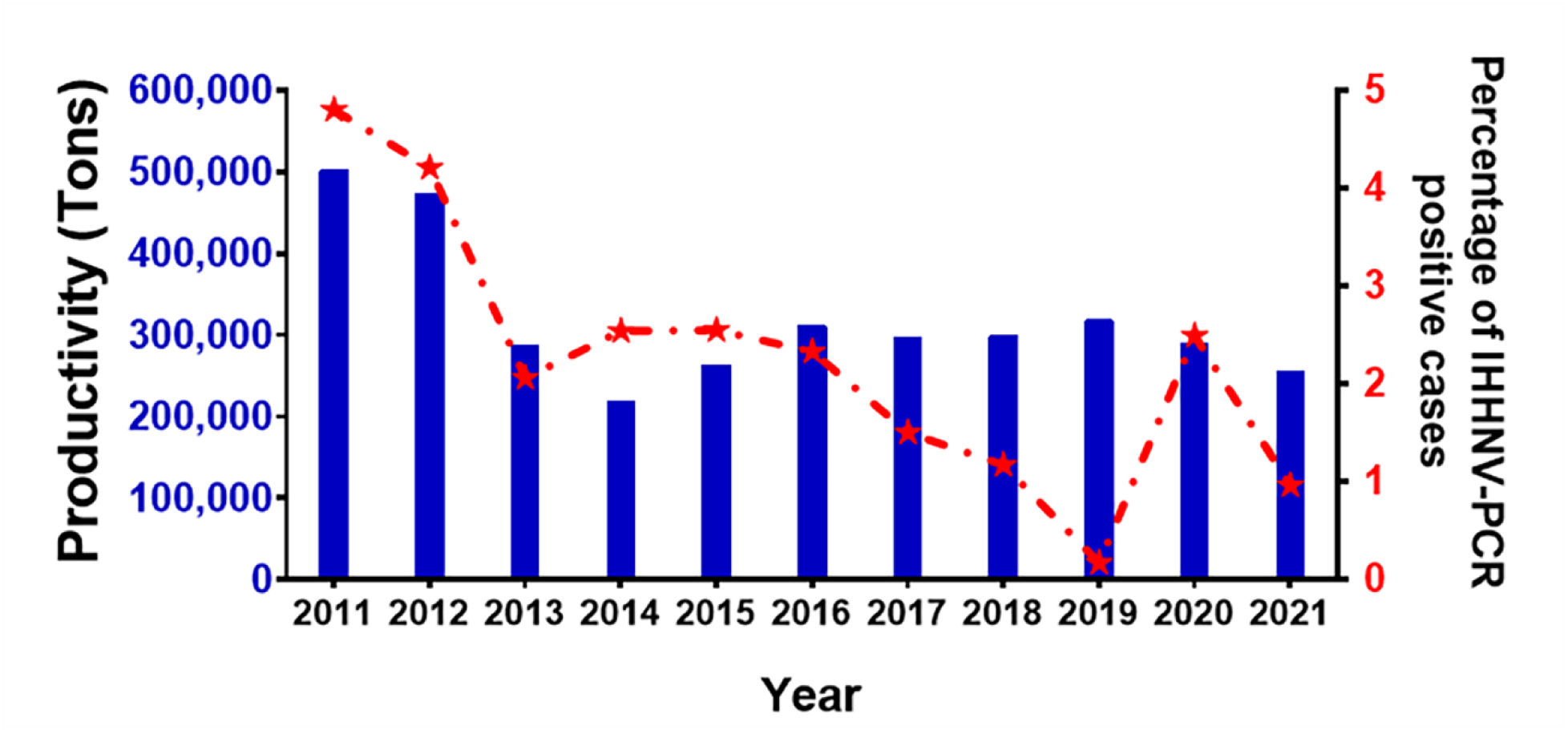
Thailand shrimp production (blue bars and left Y axis) for both *P*. *monodon* and *P*. *vannamei* versus the percentage of IHHNV-PCR positive cases (dotted red line and right Y axis) from 2011 to 2021. The IHHNV detection from 2011-2017 was recorded through a Thai DOF active surveillance program, whereas those in 2018-2021 were recorded through a Thai DOF passive surveillance program. The dotted line indicates the percentage (right axis) of positive IHHNV-PCR test results in the total number of samples tested for any reason by Thai DOF in each individual year.

**FIGURE 2.**
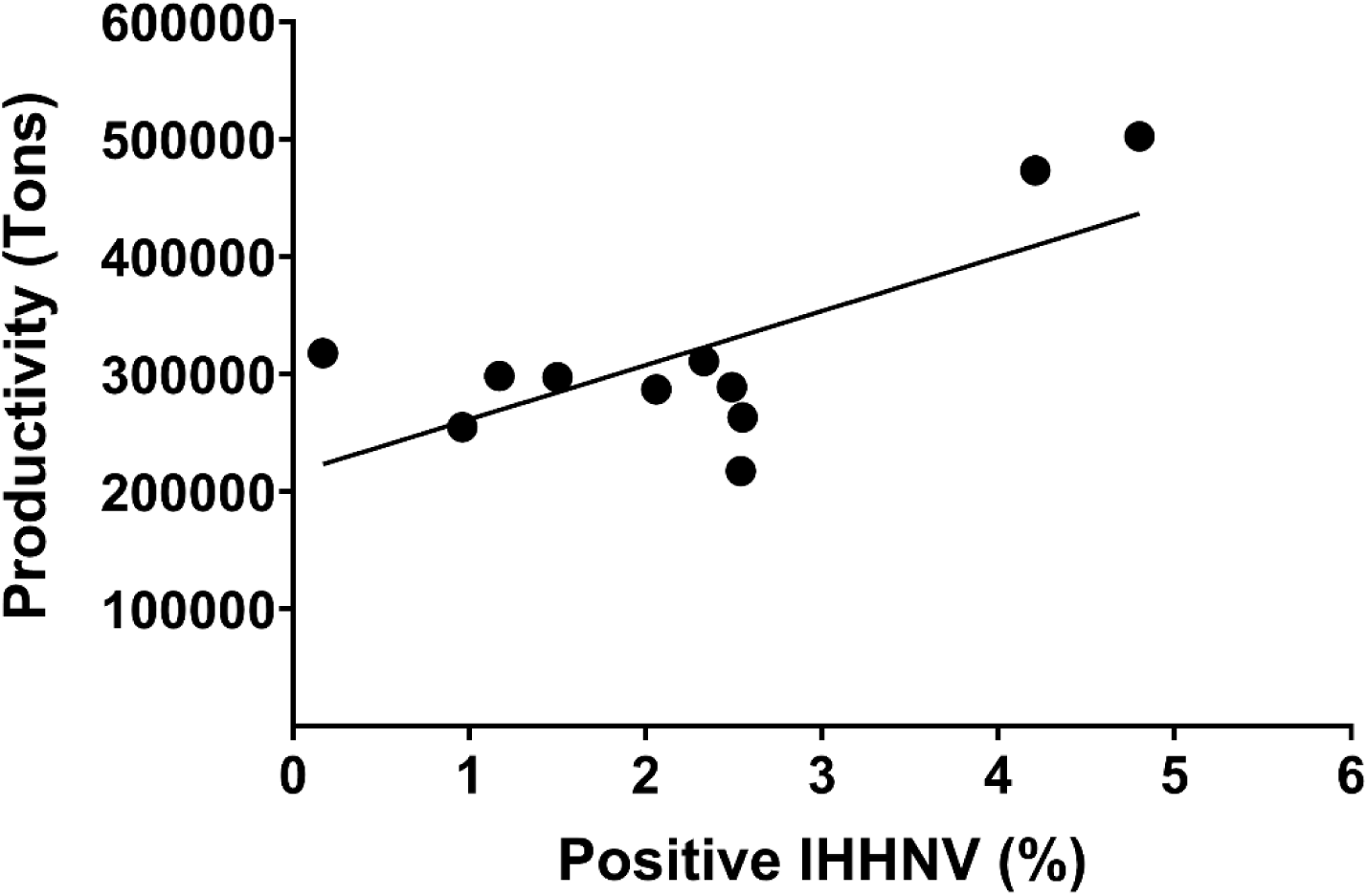
Correlation between percentage of IHHNV-PCR positive cases and shrimp production in Thailand from 2011 to 2021. The Pearson product-moment correlation coefficient (r) is 0.7 (p = 0. 015) indicating a significant positive correlation between the yearly percentage of IHHNV-PCR positive samples and yearly shrimp production.

### 3.2. Pond Sets 1 and 2 both gave profitable harvests

#### 3.2.1. Pond Set 1 production

Shrimp from all 4 culture ponds in Pond Set 1 (selected due to positive PCR test results for IHHNV) were profitably harvested with total culture periods between 123 (∼4 months) and 185 days (∼6 months) (**Table 2**). Cultivation was carried out during the months of February to August, which is the optimum period for shrimp cultivation in Thailand. The coefficients of variation for weight were in the normal range for Ponds 2 to 4 (16-25%). Only Pond #1 had an unusually high CV of 34%. However, this was because there was a long tail of very large shrimp in the groups of 36-42 g and 43-49 g that can be seen in the size distribution graph in Fig. 3. This is the opposite to RDS with CV’s at 35-90% shifted towards small sizes. Furthermore, the average daily weight gain for all the ponds in Set 1 was approximately 0.2 g, which is also normal for *P. monodon* in Thailand (Limsuwan and Chanratchakool, 2004). The farmers would not reveal their total shrimp yields or selling price, but did inform the Fisheries Department that their crops were profitable (i.e., that they did not suffer any financial losses). The time of final harvest, and sometimes partial harvesting times chosen by farmers, are dependent on highly variable market conditions. Despite these variables, the overall ADGs, shrimp sizes and CVs for ponds in Set 1 can be considered normal to better than normal for Thai conditions, and not characteristic of RDS which is characterized by a mean weight at harvest of non-profitable shrimp sizes and a CV of 35% or more shifted towards sizes of low value (Kalagayan et al., 1991).

**FIGURE 3.**
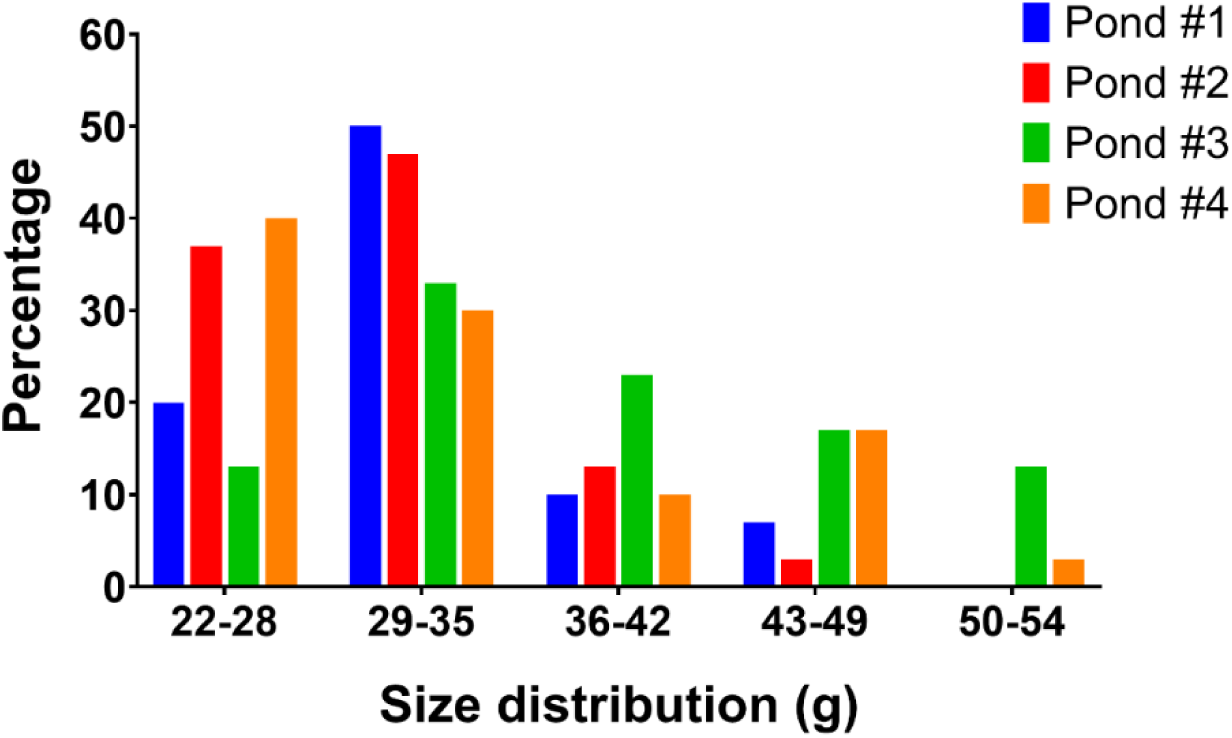
Pond Set 1 bar graph showing the size distribution of the shrimp in the 4 ponds studied in Set 1. The medians for all the ponds were in the range of 22 to 35g.

**TABLE 2.** Pond Set 1 shrimp growth performance and pond productivity.

| Pond | Stocking density (shrimp/m<sup>2</sup>) | Duration of culture (days) | CV* (%) | Mean harvested weight (g) (mean± SD) | ADG (g) |
| --- | --- | --- | --- | --- | --- |
| 1 | 36 | 123 | 34 | 25.2±8.50 | 0.20 |
| 2 | 50 | 152 | 19 | 30.9±5.97 | 0.20 |
| 3 | 54 | 149 | 22 | 37.0±8.13 | 0.25 |
| 4 | 55 | 185 | 25 | 32.5±8.14 | 0.18 |

Note: CV: Coefficient of Variance; ADG: Average Daily weight Gain.

#### 3.2.2 Pond Set 2 production

Similar to Pond Set 1, all ponds in Set 2 gave profitable harvests (**Table 3**) with cultivation times ranging from 120 days (∼4 months) to 174 days (∼6 months). This included Ponds #1 and #7 that showed abnormally high CV (41% and 39%, respectively). However, the mean size for Pond #1 was 29±12 g and the median (**Figure 4**, blue arrow) was 36 to 42 g. This indicated that the high CV was due to a wide range of sizes shifted towards large sizes (similar to Pond 1 in Set 1, and a profile opposite to the RDS profile). On the other hand, Pond #7 did have the smallest mean size (17±7 g) and smallest median size (15-21 g) (**Figure 4**, gray arrow) in pond Set 2. Thus, the size distribution and CV (39%) were similar to those of an RDS pond. Pond #6 (**Figure 4**, black bar) was intermediate, with a CV of 24%, a mean size 20±5 g and a median size of 15-21 g (**Figure 4**, black arrow). Like Pond #1, this does not fit the RDS profile. In summary, only Pond #7 showed a CV (39%) and size profile that corresponded with the description of RDS. However, this was only in terms of size distribution (i.e., there was no shrimp deformity). Despite the differences among these ponds and despite the profile of Pond #7, all the farmers claimed profitable crops. This was because the cultivations were carried out during the months of October to April which is the riskiest “Thai winter season” characterized by low temperatures conducive to outbreaks of white spot disease (WSD) caused by white spot syndrome virus (WSSV). However, during this period shrimp prices are high, attracting some farmers to attempt cultivation despite the risks.

**FIGURE 4.**
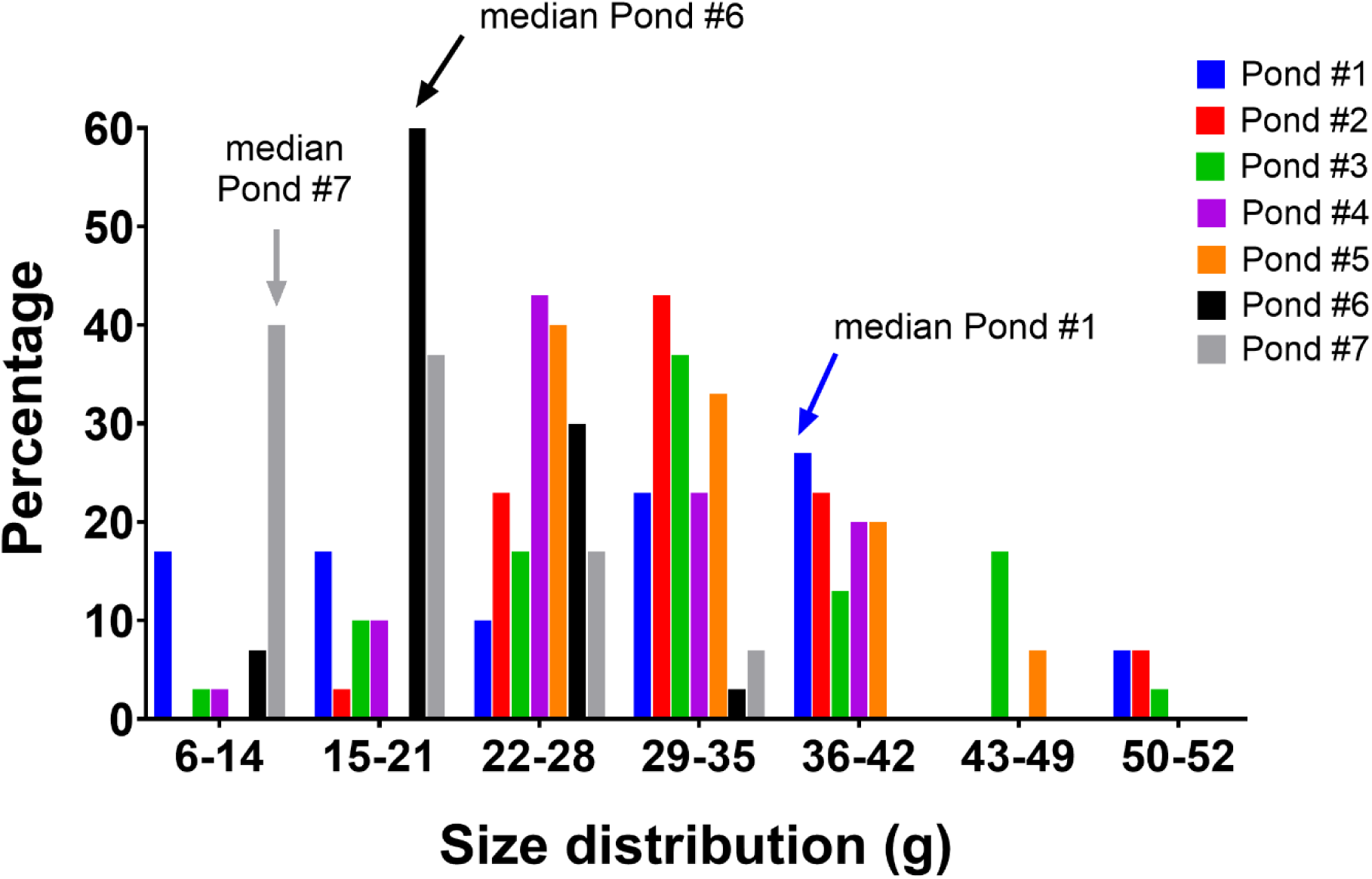
Pond Set 2 bar graph showing the shrimp size distribution in the 7 ponds studied.

**TABLE 3.**
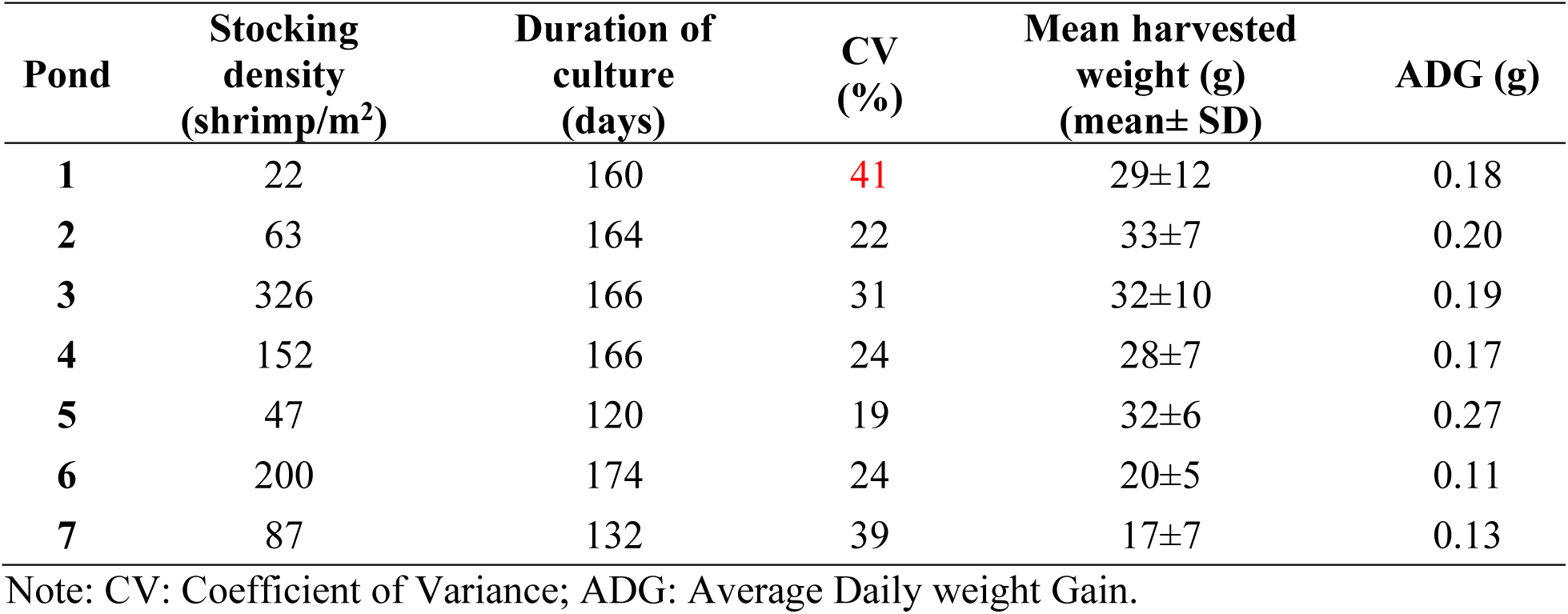
Pond Set 2 shrimp growth performance. Note: CV: Coefficient of Variance; ADG: Average Daily weight Gain.

| Pond | Stocking density (shrimp/m<sup>2</sup>) | Duration of culture (days) | CV (%) | Mean harvested weight (g) (mean± SD) | ADG (g) |
| --- | --- | --- | --- | --- | --- |
| 1 | 22 | 160 | 41 | 29±12 | 0.18 |
| 2 | 63 | 164 | 22 | 33±7 | 0.20 |
| 3 | 326 | 166 | 31 | 32±10 | 0.19 |
| 4 | 152 | 166 | 24 | 28±7 | 0.17 |
| 5 | 47 | 120 | 19 | 32±6 | 0.27 |
| 6 | 200 | 174 | 24 | 20±5 | 0.11 |
| 7 | 87 | 132 | 39 | 17±7 | 0.13 |

Note: CV: Coefficient of Variance; ADG: Average Daily weight Gain.

### 3.3. High frequency of false positive IHHNV PCR test results with WOAH methods

#### 3.3.1. Pond Set 1 IHHNV PCR test results

Using a plasmid carrying the target sequence for IHHNV, the detection sensitivity of the IHHNV-LA nested PCR method was found to be 10 copies (**Supplementary Figure S1**). Of the DNA extracts from 5 shrimp randomly collected from each of the 4 ponds in Pond Set 1, all gave positive PCR test results with both the OIE-IHHNV-309 and OIE-IHHNV-389 methods, while only 2 ponds (Pond #1 and #2) gave additional positive PCR test results with the IHHNV-LA method. Representative agarose gels of PCR amplicons from Pond Set 1 are shown in **Figure 5** and the summarized test results for Pond 1 are shown in **Table 4**. The negative PCR test results obtained using the LA method raised the possibility that 10 specimens in Ponds 3 and 4 had given false positive test results for IHHNV using the 2 WOAH methods.

**FIGURE 5.**
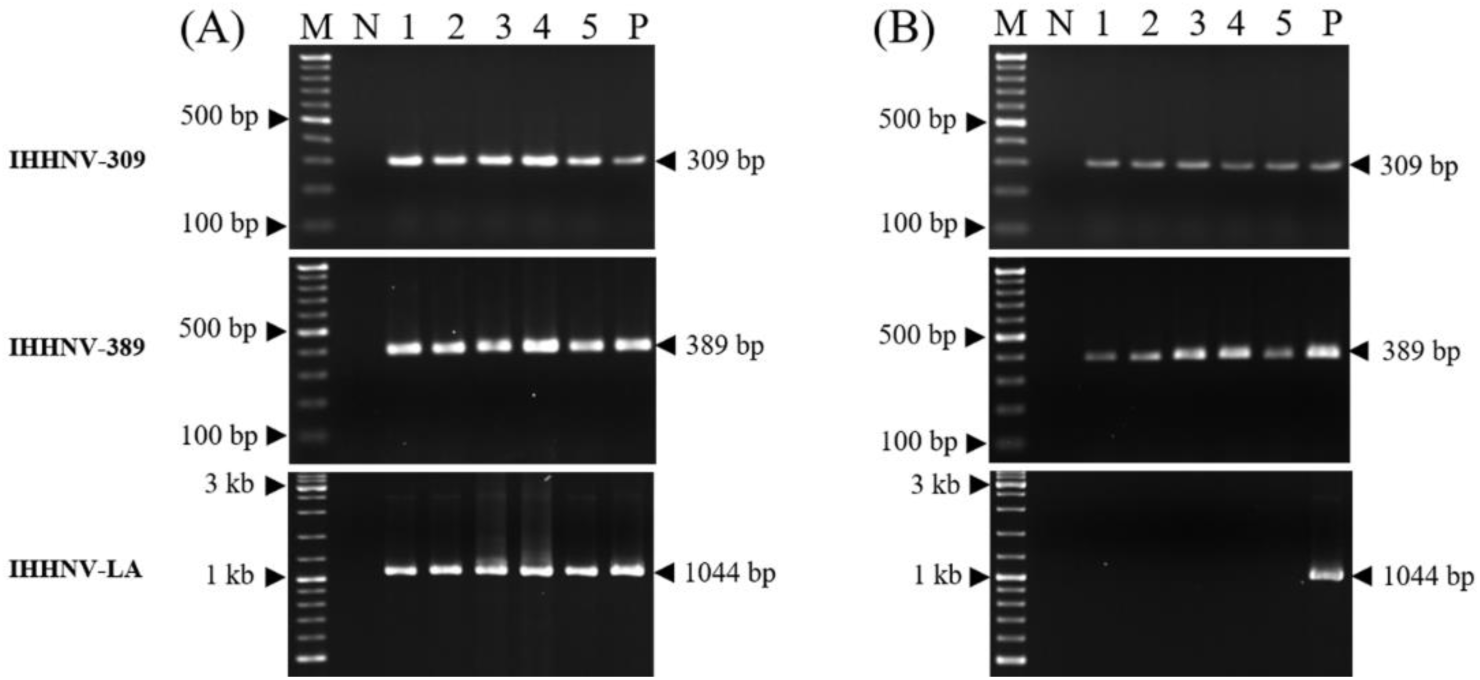
Example gels of PCR amplicons from shrimp specimens using 3 IHHNV-PCR detection methods. (A) Agarose gels showing positive PCR test results for all 3 PCR detection methods with 5 representative specimens from Pond #1. (B) Agarose gels from representative specimens from Pond #3 showing positive PCR test results with the 2 OIE methods only and negative results for the IHHNV-LA method.

**TABLE 4.** Pond Set 1 summary of IHHNV-DNA detection by PCR in juvenile shrimp from 4 culture ponds. Gray background highlights ponds positive by all 3 PCR methods.

| Pond<br># | PCR methods |  |  |
| --- | --- | --- | --- |
|  | IHHNV-309 | IHHNV-389 | IHHNV-LA |
| 1 | 5/5 | 5/5 | 5/5 |
| 2 | 5/5 | 5/5 | 5/5 |
| 3 | 5/5 | 5/5 | 0/5 |
| 4 | 5/5 | 5/5 | 0/5 |

Notes: LA: Long-Amp PCR

#### 3.3.2. Pond Set 2 IHHNV PCR test results

The 6 shrimp samples individually tested for IHHNV from each of 7 ponds in Set 2 gave positive test results with both WOAH-recommended methods (**Table 5**). However, with the IHHNV-LA method, only specimens from Pond #3 (1 in 6) and Pond #7 (3 in 6) gave positive PCR test results. This indicated that 5 out of 7 ponds had given false positive PCR test results for infectious IHHNV using the 2 WOAH methods. For Ponds #3 and #7, the results suggested that the prevalence of IHHNV was higher in Pond #7 than Pond #3.

**TABLE 5.** Pond Set 2 summary of IHHNV-DNA detection by PCR. Gray background highlights the two ponds giving positive results with all 3 PCR test methods.

| Pond<br># | PCR methods for IHHNV |  |  |
| --- | --- | --- | --- |
|  | IHHNV-309 | IHHNV-389 | IHHNV-LA |
| 1 | 6/6 | 6/6 | 0/6 |
| 2 | 6/6 | 6/6 | 0/6 |
| 3 | 6/6 | 6/6 | 1/6 |
| 4 | 6/6 | 6/6 | 0/6 |
| 5 | 6/6 | 6/6 | 0/6 |
| 6 | 6/6 | 6/6 | 0/6 |
| 7 | 6/6 | 6/6 | 3/6 |

### 3.4. Cowdry A type inclusion (CAI) analysis did not match IHHNV PCR results

#### 3.4.1. Pond Set 1 IHHNV infection confirmed by CAI only in Pond #1

Pond Set 1 specimens from all 4 ponds (5 each, total 20) were subjected to histological examination for CAI. An example of IHHNV-CAI is shown in **Figure 6a** from a slide of archived positive control tissue. From Pond Set 1, only 1 specimen from Pond #1 showed 1 CAI in the antennal gland (**Table 6**; **Figure 6c**) and 2 in the nerve cord (**Figure 6d**). There were no positive cells of the hematopoietic tissue, the cuticular epidermis or any other IHHNV target tissues. In other words, only 1 of 5 specimens from 1 pond positive for IHHNV with all three PCR methods showed CAI, and even then, at a very low prevalence. In contrast, **Figure 6b** shows an example of normal hematopoietic tissue (HPT) seen in samples from the study ponds.

**FIGURE 6.**
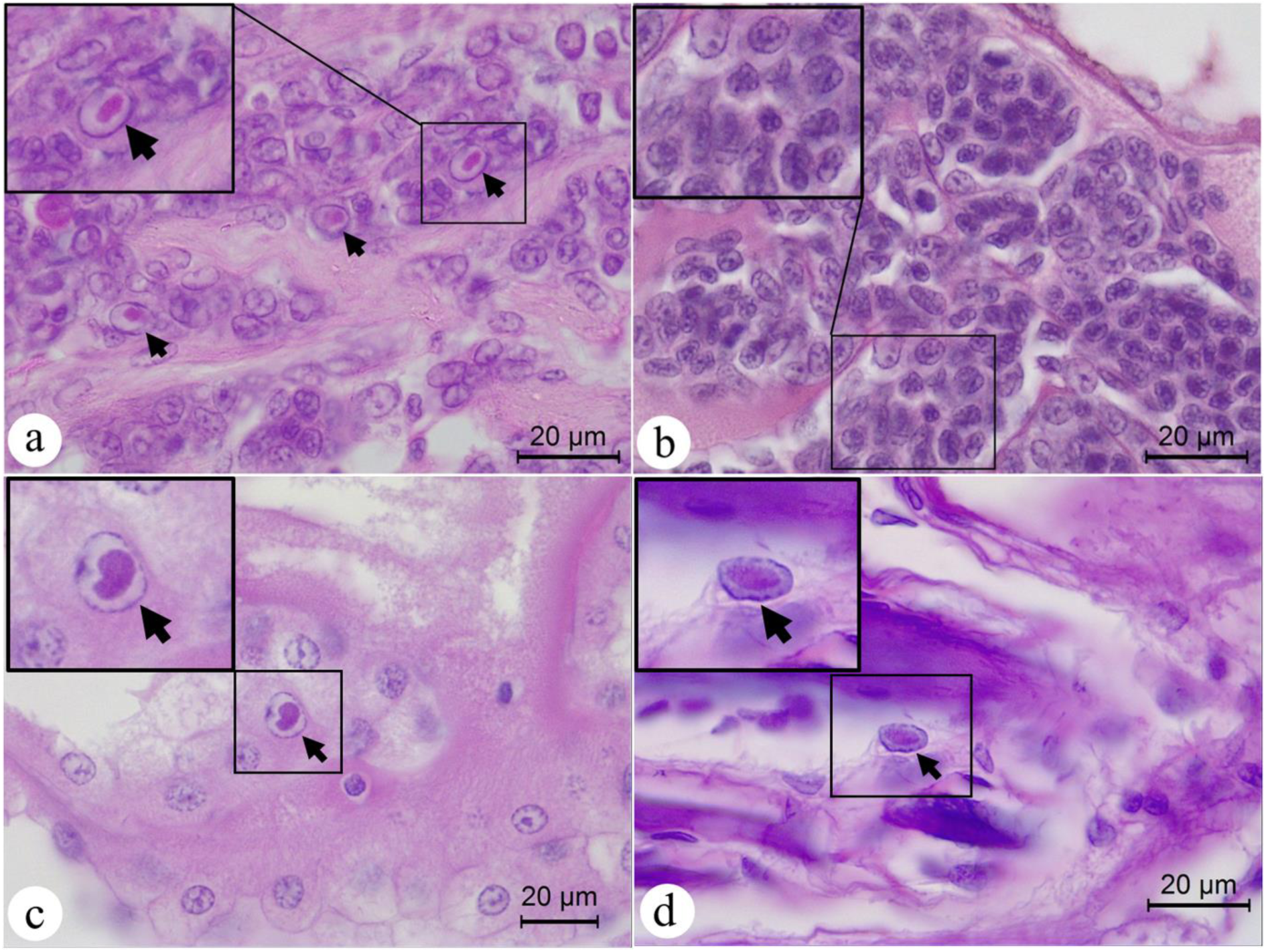
Example photomicrographs of tissues positive or negative for IHHNV by normal histology. **(a)** Example photomicrograph from an archived positive-control specimen showing IHHNV characteristic lesions (Cowdry Type A inclusions or CAI) (black arrows), with the upper right inclusion magnified in the insert at the upper left. **(b)** Example photomicrograph of normal HPT (i.e., no CAI) from one of the study specimens positive for IHHNV using the IHHNV-LA method. **(c, d)** Photomicrographs of the antennal gland and nerve cord showing Cowdry Type A inclusions or CAI, respectively, from a single specimen (Pm 1.1) from Pond Set 1.

**Table 6.**
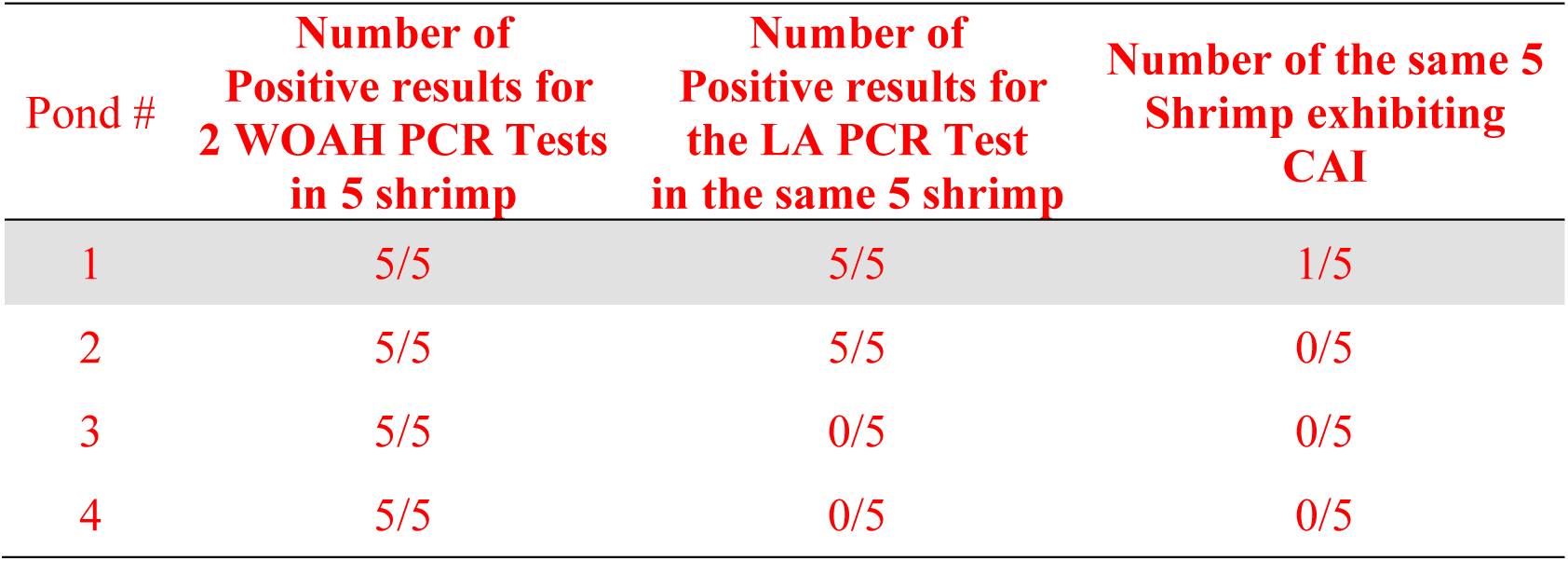
Pond Set 1 results of histological analysis for CAI in shrimp. Note that only 1 specimen out of 5 from Pond 1 (gray background) gave positive test results for CAI.

#### 3.4.2. Pond Set 2 IHHNV infection confirmed by CAI only in Pond 7

Out of the 7 ponds in Set 2, only Ponds #3 and #7 gave positive test results for IHHNV using all 3 PCR methods. As stated in the M&M section, the 6 specimens from each pond in Set 2 used for PCR testing were different from the 3 specimens from each pond prepared for histological analysis via CAI, ISH and IHC. Results in **Table 7**. show that 2 of 3 specimens from Pond #7 only exhibited the presence of CAI. It is possible that, that Pond #3 may have yielded some positive specimens if more samples had been provided.

**TABLE 7.** Pond Set 2 summary of IHHNV-DNA detection by PCR and by presence of CAI. Note that only 2 of 3 specimens from Pond #7 (gray background) gave positive test results for IHHNV by the presence of CAI.

| Pond # | Number of positive results for the LA-PCR Test in 6 shrimp | Number exhibiting CAI in 3 different Shrimp |
| --- | --- | --- |
| 1 | 0/6 | 0/3 |
| 2 | 0/6 | 0/3 |
| 3 | 1/6 | 0/3 |
| 4 | 0/6 | 0/3 |
| 5 | 0/6 | 0/3 |
| 5 | 0/6 | 0/3 |
| 7 | 3/6 | 2/3 |

### 3.5. ISH test results matched CAI results for both Pont Sets

#### 3.5.1. Pond Set 1 ISH-positive results matched CAI results

For ISH tests using probes based on amplicons from the two WOAH detection methods (IHHNV-309 and IHHNV-389), only 2 ponds were tested with both probes. These were Pond #1 that had given positive results for CAI and Pond #3 that had given negative results for CAI (**Table 8**, **Figure 7-8**). **Figure 7** shows representative photomicrographs. In Fig. 7a-c, the positive control specimen shows strong ISH reactions (dark staining) in the HPT. Specimen Pm 1.1 from Pond Set 1 that did show CAI gave positive ISH test results with both probes in the HPT (Fig. 7d, e) and the Cep (Fig. 7g, h). In contrast, Pm 1.3 that did not show CAI gave negative ISH results with both probes in both the HPT (Fig. 7j, k) and the cuticular epithelium (Cep) (Fig. 7m, n). These ISH results corresponded with the CAI results, but not with the LA-PCR results. Although the proportional numbers of IHHNV positive cells in the HPT were much lower than in the Cep in Fig. 7, highly variable loads of IHHNV in different tissues of the same shrimp specimen and between shrimp specimens have been previously reported for IHHNV (Chayaburakul et al., 2005).

**Figure 7.**
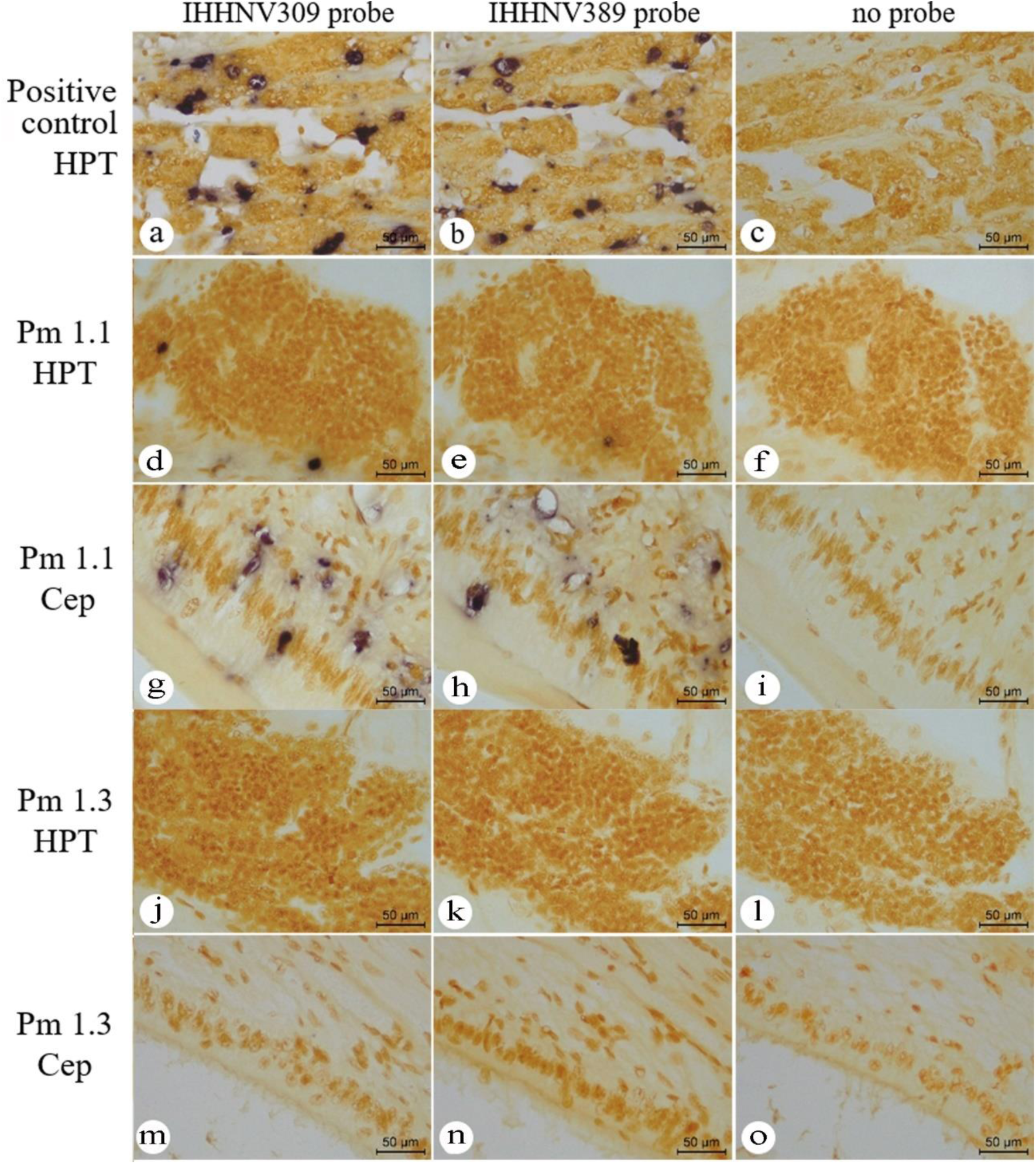
Pond Set 1, representative photomicrographs of ISH test results from adjacent tissue sections for 2 specimens (1.1 and 1.3) from Pond #1. (a,b) Photomicrographs showing positive ISH results (dark staining) with both probes using the positive control for IHHNV. (c,f,i,l,o) This column on the right shows the results for the no-probe negative control. (d,e,g,h) Positive ISH results for specimen Pm 1.1 with both probes in both the HPT and Cep., (j,k,m,n) Negative ISH reactions with both probes in HPT and Cep with specimen 1.3.

**Figure 8.**
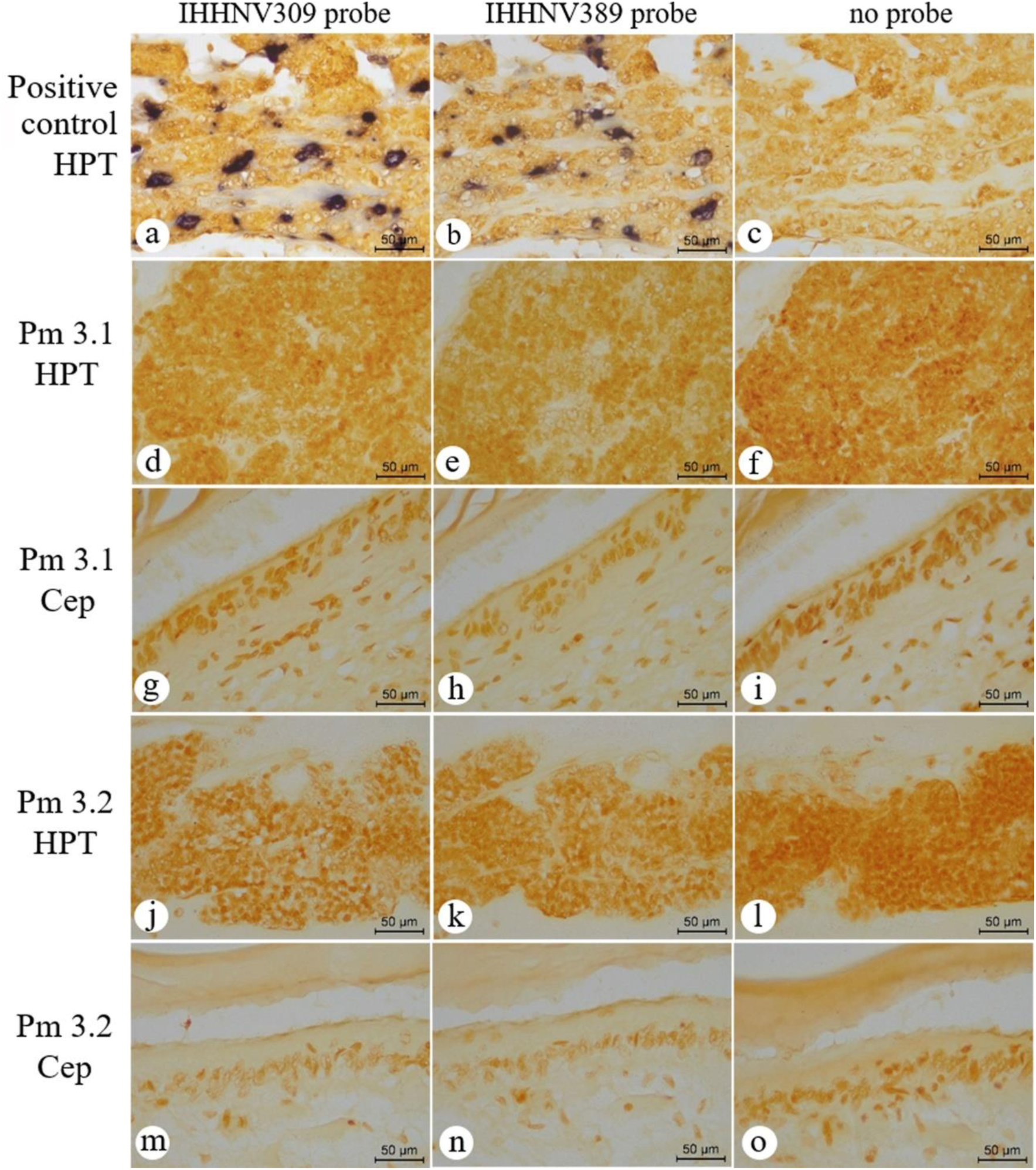
Pond Set 1, representative photomicrographs of ISH test results from adjacent tissue sections for 2 specimens (Pm 3.1 and 3.2) from Pond #3. **(a,b)** Photomicrographs showing positive ISH results (dark staining) for IHHNV with the same positive control as in Fig. 6. The column on the right **(c,f,i,l,o)** shows the no-probe negative controls. There were no positive ISH results in either of the two specimens with both probes and both tissues **(d,e,g,h,j,k,m,n)**.

**TABLE 8.** Pond Set 1 agreement between ISH and CAI results for ponds 1 and 3. The positive pond is highlighted in gray background.

| Pond # | Number of Positive LA PCR results in 5 shrimp | Number of Positive results for CAI in same 5 shrimp | Number of Positive results for ISH in same 5 shrimp |
| --- | --- | --- | --- |
| 1 | 5/5 | 1/5 | 1/5 |
| 3 | 0/5 | 0/5 | 0/5 |

#### 3.5.2. Pond Set 2 ISH-positive results also matched CAI-positive results

Out of the 7 ponds in Set 2, only 2 specimens in 3 (Pm Pond #7) gave positive test results for IHHNV by ISH and presence of CAI in 2 of 3 specimens tested (**Table 9**). It is possible that if more samples had been provided, that Pond #3 may have yielded some positive specimens. In any case, the CAI result and ISH results match with respect to pond status for IHHNV presence. In addition, for Pond #7, the result for only 2 ISH positive suggests that the ISH test is less sensitive than PCR or that the LA-PCR test might give false positive test results for IHHNV.

**Table 9.** Pond Set 2 summary of positive ISH results (gray background) compared with LA-PCR and CAI test results.

| Pond # | Number of positive results for the LA-PCR Test in 6 shrimp | Number exhibiting CAI in 3 different Shrimp | Number of Positive results for ISH in the 3 shrimp |
| --- | --- | --- | --- |
| 1 | 0/6 | 0/3 | 0/3 |
| 2 | 0/6 | 0/3 | 0/3 |
| 3 | 1/6 | 0/3 | 0/3 |
| 4 | 0/6 | 0/3 | 0/3 |
| 5 | 0/6 | 0/3 | 0/3 |
| 5 | 0/6 | 0/3 | 0/3 |
| 7 | 3/6 | 2/3 | 2/3 |

### 3.6. IHC test results also matched CAI and ISH results for both Pond Sets

#### 3.6.1. Pond Set 1 IHC results

The same 5 shrimp specimens used for ISH tests were also subjected to IHC analysis using an anti-IHHNV capsid protein antibody. Results obtained using the IHC method (**Table 10**) exactly matched the positive and negative ISH results obtained using those same specimens **(Table 8**). Specifically, the positive results (brown staining) and negative results (no brown staining) for each of the specimens matched the positive and negative ISH results for the same shrimp specimens (**Figures 9** and **10)**. Only one specimen, Pm 1.1 from Pond #1, gave positive IHC results (**Figure 9**), corresponding with the CAI and ISH positive results and with the PCR positive results by all 3 PCR methods. In contrast Pm 1.3 from Pond #1 and Pm 3.1 and 3.2 from Pond #3 all gave negative IHC test results. As with the ISH tests, these IHC results indicated that even the IHHNV-LA method may give false positive test results for the presence of infectious IHHNV.

**FIGURE 9.**
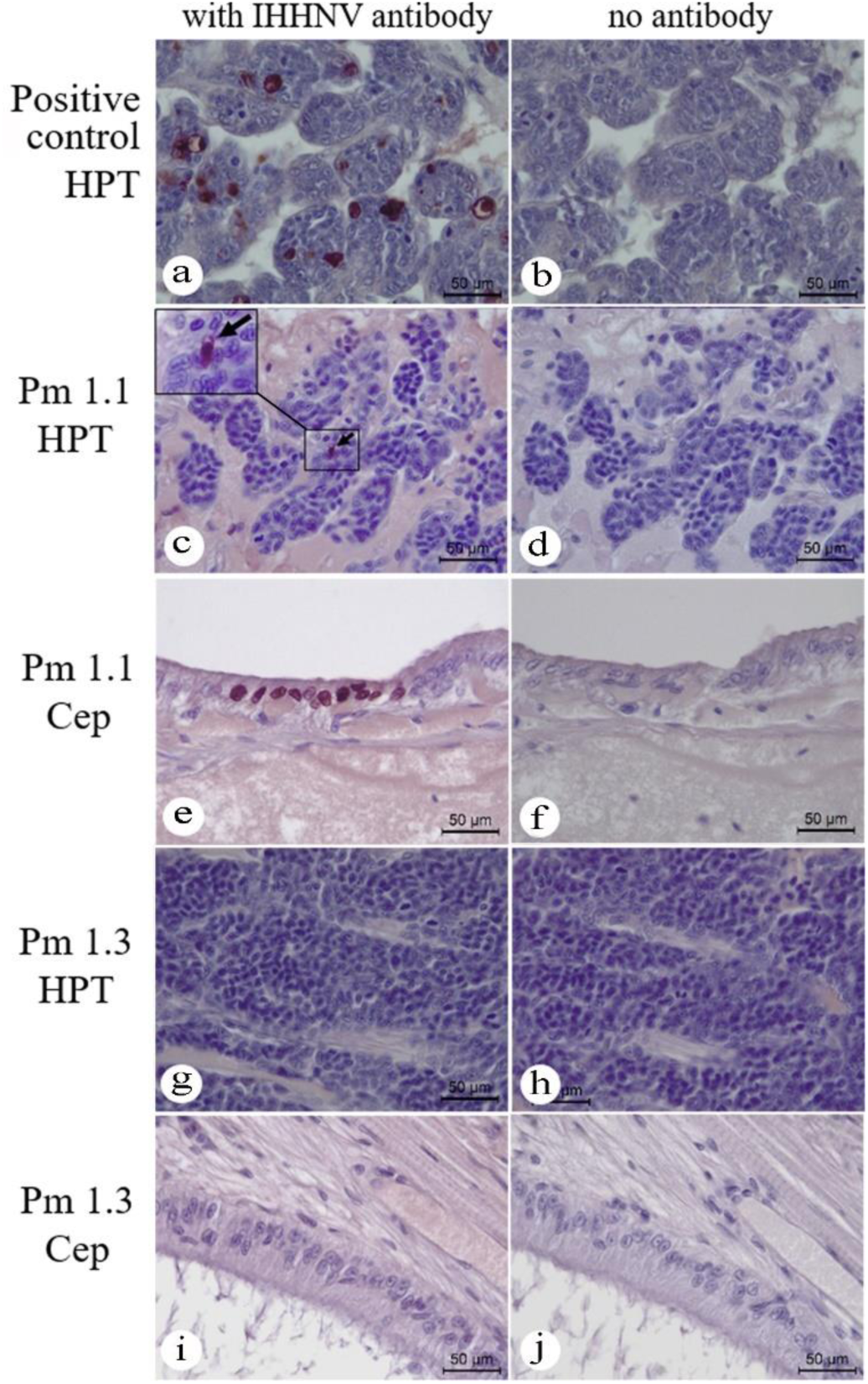
Pond Set 1 representative photomicrographs of immunohistochemistry (IHC) results for the same 2 shrimp specimens (Pm 1.1 and Pm 1.3) from Pond #1 used for ISH experiments. (a, b) Archived positive control shrimp tissue infected with IHHNV and showing positive IHC staining (dark brown) in the HPT. (c, d) Positive IHC results for HPT of Pm 1.1. (e, f) Positive IHC results for Cep of Pm 1.1. (g, h) Negative IHC results for HPT of Pm 1.3. (i, j) Negative IHC results for Cep of Pm 1.3.

**FIGURE 10.**
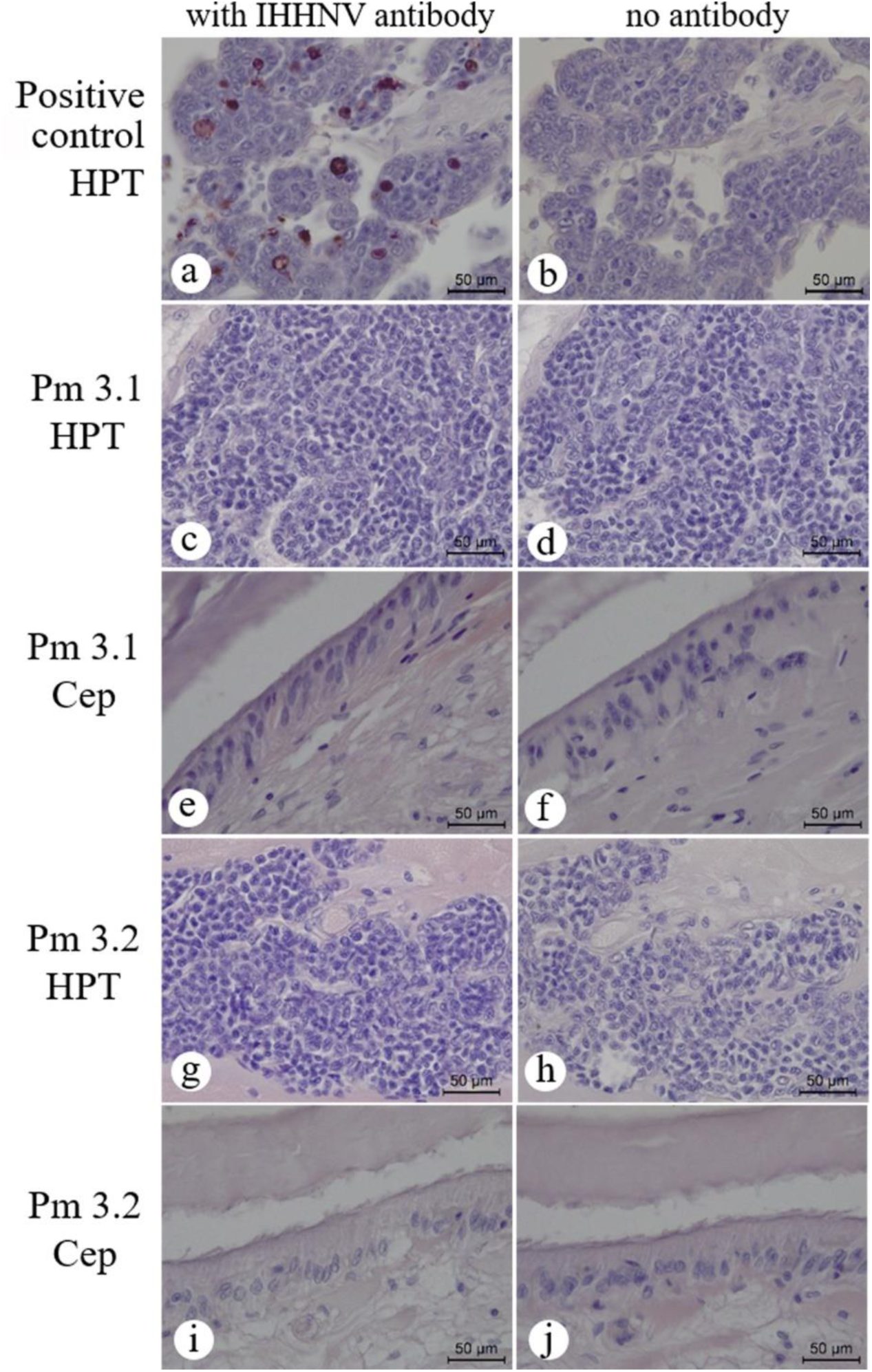
Pond Set 1 representative photomicrographs of immunohistochemistry (IHC) results. Example photomicrographs of IHC results from the same two specimens (Pm 3.1 and Pm 3.2) from Pond #3. (a, b) Archived positive control shrimp infected with IHHNV showing positive IHC staining (dark brown) in the HPT. All other photomicrographs for both tissues show negative IHC results (no brown staining).

**Table 10.** Pond Set #1 IHC test results matched CAI and ISH results.

| Pond # | Number of<br>Positive LA-PCR<br>results in 5 shrimp | Number of<br>Positive results for<br>CAI in same 5 shrimp | Number of<br>Positive results for<br>IHC in same 5 shrimp |
| --- | --- | --- | --- |
| 1 | 5/5 | 1/5 | 1/5 |
| 3 | 0/5 | 0/5 | 0/5 |

#### 3.6.2. Pond Set 2 IHC results matched CAI and IHC results

Like Pond Set #1, the same 3 shrimp specimens from Pond Set #2 previously used for CAI detection and ISH testing were used for IHC testing and gave results shown in **Table 11**. Only Pond 7 gave positive IHC results for the same 2 specimens that exhibited CAI.

**Table 11.** Pond Set 2 comparison of IHC test results with CAI and ISH results. Only samples from Pond #7 gave positive test results (gray background).

| Pond # | Number of 3 shrimp exhibiting CAI | Number of 3 shrimp positive by ISH | Number of 3 Shrimp positive by IHC |
| --- | --- | --- | --- |
| 1 | 0/3 | 0/3 | 0/3 |
| 2 | 0/3 | 0/3 | 0/3 |
| 3 | 0/3 | 0/3 | 0/3 |
| 4 | 0/3 | 0/3 | 0/3 |
| 5 | 0/3 | 0/3 | 0/3 |
| 5 | 0/3 | 0/3 | 0/3 |
| 7 | 2/3 | 2/3 | 2/3 |

### 3.7. Pathogens other than IHHNV were also present in both pond sets

#### 3.7.1. Pathogens other than IHHNV found in Pond Set 1

In addition to microscopic examination for CAI, all 20 shrimp specimens in Pond Set 1 were examined for other histopathology that might be associated with retarded growth (e.g., MBV, HPV, EHP and bacteria in the HP). No signs of MBV, HPV or EHP were seen in the HP of any of the 20 specimens. However, some specimens showed the presence of bacterial granulomas, some of which were melanized, indicating long presence. Other anomalies seen are summarized in **Table 12**.

**TABLE 12.**
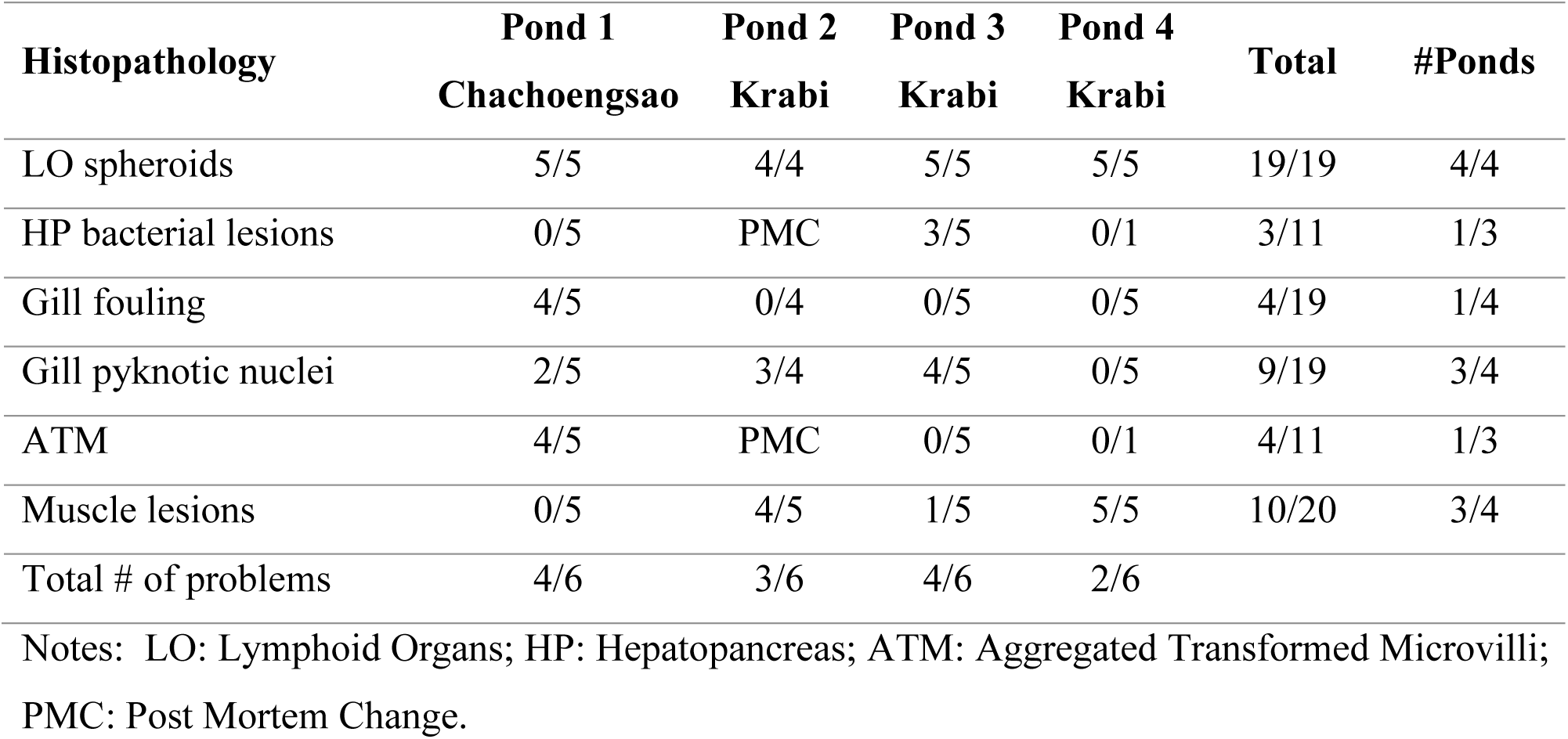
Other histopathological signs in tissues of the 20 specimens from the 4 study ponds. Totals for each pond less than 5 indicate absence of the target tissue in the microscope slide. PMC indicates postmortem change from poor tissue fixation of the target organ, which makes it difficult to assess pathology. Example photomicrographs of the various histopathology’s are shown in **Supplementary Figure 2.**

None of the anomalies (e.g., LO spheroids, HP lesions, etc.) listed in Table 4 gave ISH-positive or IHC-positive signals using the IHHNV-specific probes. Thus, absence of ISH and IHC signals indicated that the histological anomalies did not contain detectable IHHNV. Nor have such anomalies been reported to be caused by IHHNV. Lymphoid organ (LO) spheroids develop in response to foreign material accumulated by phagocytic hemocytes and also during viral infections with Taura syndrome virus (TSV) (Hasson et al., 1999) and yellow head virus (YHV) (Soowannayan et al., 2003). The spheroids caused by these pathogens are positive by ISH and IHC specific for the respective viruses. Thus, we can assume that the spheroids in our specimens were not formed in response to IHHNV but to other causes.

The other problems exhibited in the specimens were the presence of bacterial lesions in the HP, pyknotic nuclei in gill tissue, muscle necrosis and gill fouling. According to the Lightner scale of disease severity (Lightner, 1996), these other problems were present at the levels from “Trace” (Signs of infection/infestation by internal pathogens, parasite or epicommensals present at just above diagnostic procedure minimum detection limits) or “1” (“Signs of infection/infestation by pathogen, parasite or epicommensal present, but at levels that may be below those needed for clinical disease”). These levels matched those for IHHNV in the few specimens where IHHNV-CAI were observed.

The presence of LO spheroids in specimens from all ponds indicated a substantial immune response to an unknown factor in all ponds and this was compounded by additional sub-clinical problems that varied from pond to pond, also from undetermined causes. In addition, gill fouling is a sign of low pond water quality leading to reduced ability of shrimp to maintain clean gills, and muscle necrosis is a sign of shrimp stress. It is important that even 1 positive specimen for any pathogen in a random sample of 10 specimens is indicative of 26% or more prevalence with 95% confidence in the source population (Cameron, 2002). Thus, even 1 positive sample in 6 or 5 specimens would indicate prevalences of 40 to 46% or more. As a result, the histological information clearly indicates that conclusions about the effect of IHHNV on production from these ponds based solely on PCR detection of IHHNV would be scientifically untenable.

#### 3.7.2. Pathogens other than IHHNV in Pond Set 2

The 3 specimens examined by PCR, ISH and IHC in Pond Set 2 (**Table 7**) were also examined histologically in H&E-stained tissue sections for the presence of pathologies other than IHHNV (**Table 13**). Because of the small number of shrimp sampled, even 1 positive specimen in 3 indicated a very high prevalence (64% or more) in the source population. Similar to Pond Set 1, all samples from every pond (total 3 x 7 = 21) showed the presence of lymphoid organ spheroids indicative of an immune response by phagocytic cells (Hasson et al., 1999) but not a response to IHHNV because of the negative ISH and IHC reactions in the LO to both types of IHHNV probes.

**Table 13.** Other histopathological signs in shrimp tissues from 7 ponds of Pond Set 2. Note that samples (grey background) from Ponds 1, 2 and 3 gave high estimated prevalence for HPV (gray background) and from Pond 7 gave a high estimated prevalence for MBV (gray background), both of which are known to be associated with retarded growth in *P. monodon*. Example photomicrographs of the histopathologies are shown in **Supplementary Figure 3**.

| Histopathology | Pond 1<br>Phuket | Pond 2<br>Phuket | Pond 3<br>Phuket | Pond 4<br>Phuket | Pond 5<br>Phuket | Pond 6<br>Phuket | Pond 7<br>Phang<br>Nga | Total | #<br>Ponds |
| --- | --- | --- | --- | --- | --- | --- | --- | --- | --- |
| LO spheroids | 3/3 | 3/3 | 3/3 | 3/3 | 3/3 | 3/3 | 3/3 | 19/21 | 7/7 |
| HP bacterial lesions | 0/3 | 0/3 | 0/3 | 1/3 | 0/3 | 1/3 | 1/3 | 3/21 | 3/7 |
| HP hemocyte aggregation | 0/3 | 0/3 | 1/3 | 1/3 | 3/3 | 1/3 | 0/3 | 6/21 | 4/7 |
| HP sloughing cells | 0/3 | 0/3 | 0/3 | 0/3 | 3/3 | 0/3 | 1/3 | 4/21 | 2/7 |
| HPV | 2/3 | 1/3 | 1/3 | 0/3 | 0/3 | 0/3 | 0/3 | 3/21 | 3/7 |
| MBV | 0/3 | 0/3 | 0/3 | 0/3 | 0/3 | 0/3 | 1/3 | 1/21 | 1/7 |
| Total # problems | 2/6 | 2/6 | 3/6 | 3/6 | 3/6 | 3/6 | 4/6 |  |  |

Notes: LO: Lymphoid Organs; HP: HepatoPancreas; HPV: Hepatopancreatic parvovirus; MBV: Monodon BaculoVirus.

Presence in the HP were granulomas suggestive of bacterial infections (4/7 ponds), hemocyte aggregation (4/7 ponds) indicating an immune response, and cell sloughing (2/7 ponds) similar to that caused by acute hepatopancreatic necrosis disease (AHPND). In addition, Ponds #1, #2 and #3 gave high estimated prevalences for HPV and Pond #7 gave a high prevalence for MBV, both of which are known to be associated with retarded growth in *P. Monodon* (Flegel et al., 2004, Flegel, 2006). Pond #6 also had a median of relatively small shrimp sizes, but was negative for IHHNV by all tests and also negative for HPV and MBV, clearly showing that retarded growth can arise from causes other than IHHNV. In contrast, Pond #3 positive for IHHNV by the IHHNV-LA method might have been confirmed for IHHNV by histology if more samples had been taken, but it showed a normal to better than normal size profile. Each pond exhibited from 2 to 4 of these 6 histological anomalies. Clearly, any proposal regarding the shrimp production in these study ponds based solely on IHHNV-PCR detection results and ignorance of the presence or absence of other possible pathologies would be scientifically unsupportable. This is especially important when one considers that HPV and MBV are both known to cause retarded growth in *P. monodon* (Flegel et al., 2004, Flegel, 2006). Overall, the histological analysis of the shrimp in Pond Set 2 was worse than that in Pond Set 1 probably due to off-season cultivation.

According to the Lightner scale of disease severity (Lightner, 1996), these other problems were present at the levels from “1” (“Signs of infection/infestation by pathogen, parasite or epicommensal present, but at levels that may be below those needed for clinical disease”) to “2” (“Moderate signs of infection/infestation as shown by low to moderate numbers of parasite or epicommensal, or by number and severity of pathogen caused lesions. Prognosis is for possible production losses and/or slight increases in mortality if no treatment (if treatable) or management change is applied.”). In contrast, the IHHNV lesions (when present) were at the “Trace” level.

### 3.8. Summary of results from both pond sets

For both Pond Sets 1 and 2, any specimen giving positive test results using all three PCR test methods was not sufficient to confirm an IHHNV infection. We now believe that no conventional, single PCR method or multiple PCR method can confirm IHHNV infection. However, a positive test with any one of the three PCR methods together with a positive test with any one of the three histological methods was sufficient to confirm the presence of IHHNV infection in a pond. In addition, a positive test result with either or both WOAH PCR methods but a negative result with the IHHNV-LA method served as a negative test result for the presence of IHHNV infection. In addition to the PCR results, the histological analysis in Pond Set 1 revealed less serious pathologies than those in Pond Set 2 where additional HPV and MBV infections plus sloughing of HP tubule epithelial cells characteristic of AHPND were observed. This was probably due to off-season cultivation in Pond Set 2. Overall, the results for Pond Sets 1 and 2 were parallel with respect to IHHNV detection and lack of evidence for any direct impact by it on pond productivity.

## 4. DISCUSSION

### 4.1. IHHNV false positive PCR results frequently arise

ISH reactions for IHHNV were negative for 3/5 specimens from Pond #1 that gave PCR positive results for IHHNV using all 3 methods. It was possible that positive PCR results but negative ISH and IHC results in those 3 specimens arose because the viral loads were too low to give ISH and IHC signals. This was our former belief for shrimp positive for viral pathogens detectable only with nested PCR and negative with histochemical methods. However, the results in this study for both Pond Set 1 and Pond Set 2 revealed a very good correlation between positive ISH and IHC results with results from normal histological analysis using H&E staining to detect CAI characteristic for IHHNV. An absolute, final answer to the question of infectivity would be obtained only by preparation of purified viral extracts followed by bioassays with SPF shrimp. Lack of transmission would indicate the absence of infectious IHHNV in the extracts.

We believe it is most likely that the positive PCR results that could not be confirmed by positive results for CAI, ISH and IHC arose from endogenous viral elements (EVE) of IHHNV that contain the primer target sequences as has previously been reported (Saksmerprome et al., 2011). The presence of EVE for both IHHNV and white spot syndrome virus (WSSV) in shrimp has been confirmed (see (Flegel, 2020) for a review).

EVE with high sequence identity to infectious IHHNV were recently found (Taengchaiyaphum et al., 2022) in pseudochromosome 7 (PC7) of a published draft genome of *P. monodon* (Uengwetwanit et al., 2021). Some of these EVE showed 100% sequence identity to infectious IHHNV Type 1 with sequences that matched 100% with existing types of IHHNV at GenBank and especially with the targets of the IHHNV-309 and -389 methods. PCR tests with archived DNA from the genome project using the IHHNV-309 and -389 methods gave positive test results. However, the same DNA was negative for IHHNV using the IHHNV-LA method. Additional PCR testing confirmed that the targets for the methods were in the genomic material and not in contaminating IHHNV DNA. Thus, a DNA extract from the specimen used for sequencing would have given a positive PCR test result for infectious IHHNV, even though it was not infected.

Similar EVE in mosquitoes have been shown to be protective against their homologous viruses (Suzuki et al., 2020). Thus, eliminating EVE in breeding stocks to avoid false positive PCR test results and avoid trade restrictions might have a negative impact on farmed shrimp production if the eliminated EVE are protective. The same arguments may apply to EVE for other shrimp viruses such as TSV and WSSV that are also included in the WOAH list of controlled crustacean diseases. These precautions regarding interpretation of PCR test results have been discussed previously (Alday-Sanz et al., 2020, Saksmerprome et al., 2022).

In the end, we were forced to accept from our results herein that it would be impossible to be certain using only 1 to 3 primer sets that focus on short regions of the genome whether amplicons arose from IHHNV or EVE derived from it unless sets of 7 to 10 overlapping primers pairs were used to cover the whole IHHNV genome (Chayaburakul et al., 2005, Saksmerprome et al., 2011),. Instead, we now believe that obtaining a positive test result using any of the PCR methods recommended by WOAH (or any equivalent) must be confirmed by a histological test (CAI, ISH or IHC). One might propose that the IHHNV-LA method be used as a temporary check to screen WOAH positive specimens if histological analysis is not possible. This might reveal about half of the potential false positives and reduce unjust rejection or destruction of at least some farmed shrimp products. On the other hand, of the 10 shrimp that gave positive PCR test results with the IHHNV-LA method from Pond set 1, only 1/10 showed CAI, so there would still be a significant risk of rejections based on false positive test results. On the other hand, the use of 7 to 10 overlapping primer pairs to cover the whole IHHNV genome (Saksmerprome et al., 2011) would be an alternate choice.

### 4.2. Histopathology is required to confirm IHHNV detection by PCR

This study was carried out using shrimp (*P. monodon*) cultivation ponds that had been selected solely on the basis that sampled shrimp specimens had given positive test results using two WOAH-recommended PCR methods for detection of IHHNV infections. Such positive PCR test results are often used to justify the rejection of imported shrimp products and result in serious negative economic impacts. Here we have shown that rejections based solely on the use of either one or both of the recommended WOAH-PCR methods would be unjustified. With 11 ponds tested in the study, presence of IHHNV could be confirmed in only 2 ponds (2/11 = 18%) with 1 pond open to question (3/11 = 27%). This suggests that economically important decisions might be made based on false positive PCR test results 73 to 82% of the time.

As early as 2011 (Saksmerprome et al., 2011, Saksmerprome et al., 2022), it was revealed that fragments of the genome of existing types of IHHNV occur commonly in the shrimp genome and may collectively cover almost the whole genome (Taengchaiyaphum et al., 2022). Thus, individual specimens may potentially give false positive PCR test results for the presence of IHHNV for any target sequence. Since around 2012, such viral genome fragments have been referred to as endogenous viral elements (EVE) (Feschotte and Gilbert, 2012). Recent, analysis of the whole genome of *P. monodon* has revealed clusters of scrambled EVE from IHHNV on pseudochromosome 7 that can give false positive PCR test results for existing types of infectious IHHNV (Taengchaiyaphum et al., 2022). EVE of extant types of white spot syndrome virus (WSSV) have also been reported in *P. monodon* (Utari et al., 2017). Warnings regarding the propensity of EVE to give false negative PCR and RT-PCR test results (Saksmerprome et al., 2011, Alday-Sanz et al., 2020, Saksmerprome et al., 2022) for viral infections have been largely ignored.

Of the 3 PCR methods used in this study, none alone or in combination with the other PCR methods were sufficient to confirm the presence of IHHNV infections. Due to the existence of highly variable EVE profiles in individual shrimp specimens, confirmation required additional histological examination using H&E-stained tissues, ISH or IHC. Thus, we recommend that the WOAH diagnostic manual for shrimp viral diseases be modified to specify that no viral infection can be confirmed in shrimp by one or more positive PCR test results without additional histological analysis to demonstrate the detection of characteristic IHHNV lesions or of positive ISH or IHC results. Based on the clusters of scrambled EVE from IHHNV on pseudochromosome 7 described by (Taengchaiyaphum et al., 2022), it is possible that even using a set of 7-10 overlapping primer sets to cover the whole IHHNV genome might give a false positive test result.

### 4.3. Histopathology required to determine the true impact of IHHNV infection

In the introduction section, we reviewed the history of IHHNV infection in *P. monodon* and showed that valid reports reveal no significant negative impact on farm production (Flegel et al., 2004, Withyachumnarnkul et al., 2006). This was also the conclusion recently reported for cultivated *P. vannamei* in Ecuador and Peru (Aranguren Caro et al., 2022). For Pond Set 1 in this study, only Pond #1 contained shrimp with confirmed IHHNV infections, and its production did not differ significantly from that of the other 3 ponds in Set 1 that were negative for IHHNV. None of the ponds exhibited gross signs associated with IHHNV (e.g., RDS).

In Pond Set 2, only Pond #7 was fully confirmed to contain shrimp infected with IHHNV and the shrimp size distribution for this pond was abnormal and growth resembled that of *P. vannamei* exhibiting RDS deformities, which the *P. monodon* did not exhibit. If this pond had been tested for IHHNV by PCR only and not subjected to histological analysis, an unwary examiner might have concluded that the abnormal size distribution was caused by IHHNV. However, this conclusion would not only be contrary to the history of IHHNV in *P. monodon* but also ignorant of the presence of LO spheroids, bacterial lesions, HP cell sloughing and the presence of MBV (associated with retarded growth in *P. monodon*). The co-occurrence of these 4 pathologies in addition to IHHNV indicated a very poor overall health status for the shrimp in Pond #7. Failure to consider such issues and failure to carry out a full health analysis but instead to focus simply on PCR test results can result in flawed publications claiming negative impacts on shrimp production caused by any pathogen. Misuse of PCR detection as the single criterion in assigning IHHNV as the cause for problems with cultivated shrimp has been particularly frequent. This is most important when shrimp mortality is part of the disease expression profile since death is not a characteristic of IHHNV in shrimp other than *P. stylirostris*. Thus, Pond #7 from Pond Set 2 is an excellent example of the perils involved in using PCR detection alone to implicate IHHNV as the cause of disease expression without backup histological analysis.

### 4.4. Seedstock impacts on production must also be considered

In this study, comparison of the harvest outcomes for the shrimp ponds studied in Pond Set 1 and Pond Set 2 could not be made without considering farm management issues including details related to cultivation season, farm location and infrastructure, source of seedstock (post larvae), feed and water management, etc. Already addressed was the issue of poorer health of the shrimp in the off-season farms, but other management details were not investigated in our study. However, from Pond Set 2, the presence of HPV in Ponds #1, #2 and #3 and MBV in Pond #7 suggested that viral infections might have occurred from the surrounding environment after pond stocking. However, our previous experience has shown that these viruses are most often introduced by PL used to stock the ponds. The farms that use genuine SPF stocks free of HPV and MBV do not develop these infections in rearing ponds. Instead, unscreened PL derived from non-SPF, captured broodstock or false-SPF-derived PL from unscrupulous PL suppliers are available in Thailand at lower prices than genuine SPF-derived PL. These examples reinforce the importance for farmers to be aware of the prime needs of using SPF-derived PL and recommended biosecurity measures during cultivation.

### 4.5. The WOAH reporting for shrimp viruses needs to be modified

As stated in the Introduction, it is generally recognized in Thailand that IHHNV has had no significant negative impact on the production of cultivated *P. monodon* or *P. vannamei* since around 2010. Thus, it is not normally tested for or reported by farmers. On the other hand, samples from shrimp farms are sometimes submitted to the Thai competent authority (passive surveillance) to be tested for other reasons such as clearance before export. If such shrimp samples are tested for IHHNV by PCR for any reason and give a positive result, Thailand’s WTO obligations require that WOAH be notified, because IHHNV is a listed crustacean disease. Clearly, a report to WOAH based solely on PCR detection using the WOAH recommended method with the specimens described here for Pond #3 in Pond Set 1 would have constituted an unintended false positive report. In addition, the confirmed detection of IHHNV from Pond #1 would also be reported, but the report would not be accompanied by a note indicating that detection was not associated with any significant impact on production. This is important, because a key characteristic of a WOAH-listed disease is that it has a significant, negative economic impact on production.

This is a very important issue, given the current knowledge showing that shrimp are capable of a specific, adaptive immune response based on nucleic acids that can give rise to EVE that are heritable if they occur in germ cells (Flegel, 2020, Flegel, 2022). As a result, shrimp that survive a viral infection give rise to a vast variety of offspring, each with an individual EVE profile such that protective EVE combinations become naturally selected in nature over time or can be purposefully selected in shrimp breeding programs (Alday-Sanz et al., 2020). The eventual result is that disease outbreaks caused by the relevant pathogens decline over time as a result of genetic tolerance and development of effective prevention protocols such that the negative economic impact may decline sufficiently to justify WOAH delisting. In order to monitor this decline, it would be necessary that obligatory country reports of listed shrimp pathogens be accompanied by an estimate of any production losses associated with each report. One might propose that listing does no harm, even if there is no negative economic impact. However, this is a naïve position. For example, the cost of unnecessary PCR testing for a listed pathogen can be high and the losses arising from tests that result in unjust rejection of products can also be very high.

During the revision of this document, a new publication (Aiamsa-at et al., 2024) described a CRISPR-Cas based, isothermal detection method for IHHNV that targets its ssDNA genome only and would thus avoid interference from the dsDNA fragments of IHHNV in the shrimp genome. It is possible that such a method could be proposed for detection of IHHNV and other viruses with ssDNA genomes. On the other hand, we still maintain that making decisions regarding the impacts of viruses and other pathogens based solely on PCR results is unwise without additional information related to the source animal’s overall health status. It would be the equivalent to a physician making a medical assessment for a human patient based on a single PCR test result without including a general health examination.

## 5. CONCLUSIONS

Our research has shown that confirmation of IHHNV infection in shrimp can be confirmed by PCR detection only when combined with histopathological analysis by any 1 of 3 histological methods (i.e., H&E, ISH or IHC staining) or by H&E plus ISH or IHC. It cannot be confirmed by single or multiple PCR. There were no unprofitable crops from the sampled ponds positive for IHHNV, and only 2 of 11 ponds that gave positive PCR test results contained shrimp confirmed for IHHNV infection. These results correspond to the lack of reliable reports since 2010 on significant negative impacts of IHHNV on commercially farmed *P. monodon* or *P. vannamei* in Thailand. Similarly, there are no such reports from the world capture fishery for these shrimp species, in areas where IHHNV is known to occur. Many reports claiming losses to IHHNV focus largely or solely on PCR detection results without confirmation by histological analysis or by bioassays, and without investigation for other pathogens or without realistic evaluation of actual farm losses. Furthermore, recent reports that claim losses to IHHNV in hatcheries, experimental shrimp or on farms could have been avoided by use of IHHNV-negative broodstock and/or stocking of PL free of IHHNV combined with simple biosecurity. At the same time, we have found no reliable reports of significant losses to IHHNV from commercial cultivation ponds of the world’s major shrimp producers since 2010. A thorough investigation regarding any significant negative impact on farmed shrimp production from IHHNV is needed to reassess whether it fits the criteria required to be listed by WOAH. It is our opinion that the massive expenditure required for needless PCR testing and the current losses suffered via trade restrictions currently justified by WOAH listing are now the only significant negative economic impacts of IHHNV.

## 6. ACKNOWLEDGEMENTS

This work was funded by the National Research Council of Thailand (NRCT, grant no. N42A650869), Mahidol University under the New Discovery and Frontier Research Grant 2023 (Grant no. FF-056/2566) and the NSRF via the Program Management Unit for Human Resources & Institutional Development, Research and Innovation (Grant No: B05F640137) and the joint Chinese Academy of Science and National Science and Technology Development Agency (CAS-NSTDA) project (Grant no. P2350044). We also thank the National Science, Research and Innovative Fund, Thailand Science Research and Innovation (TSRI) (Grant no. FFB670076/0337) through the National Science and Technology Development Agency (P2351693).

## 8. ETHICS STATEMENT

This work followed Thailand’s laws for ethical animal care under the Animal for Scientific 83 Purposes ACT, B.E. 2558 (A.D. 2015) under project approval number BT-Animal document no. BT37/2563) from the National Center for Genetic Engineering and Biotechnology (BIOTEC), National Science and Technology Development Agency (NSTDA), Thailand.

## 9. AUTHOR CONTRIBUTIONS

KS, PS, JS, JT, NS, ST Methodology, Investigation, Data curation, Writing-Original draft preparation, KS, RV, CC, TWF, ST Conceptualization, supervision, manuscript writing and final approval of the manuscript.

