## Supplementary material for "No single PCR test is sufficient to determine parvovirus IHHNV presence in or impact on farmed shrimp production": 3IHHNV farm MS Supplementary information.docx

**Supplementary information 1: A detailed critique of the report of the OIE Aquatic Animal Health Standards Commission (AAHC) in September of 2020 on the delisting of IHHNV**

In the report of the OIE Aquatic Animal Health Standards Commission (AAHC) in September of 2020 in response to the request to delist IHHNV from a listed crustacean disease (Report of the meeting of the OIE Aquatic Animal Health Standards Commission (AAHC), Paris, 19‒26 February 2020 in Section 4.6, Annex 9), AAHC upheld the continued listing of IHHNV because it concluded that IHHNV fit Criterion 4b: “The disease has been shown to affect the health of cultured aquatic animals at the level of a country or a zone resulting in significant consequences e.g. production losses, morbidity or mortality at a zone or country level”. The AAHC decision report cited 22 references, 8 of which were published prior to 2010 while 14 covered the period 2010 to 2020 during which the impact of IHHNV on shrimp production should have been assessed.

Of these 14 remaining publications, 4 concerned IHHNV detection methods and 4 concerned prevalence of IHHNV by PCR detection in natural or cultured shrimp, without accompanying new data concerning farmed production impacts. Of the remaining 6 references, 1 concerned general environmental issues, 1 concerned a WSSV coinfection study and one concerned *M. rosenbergii* susceptibility to IHHNV in specimens co-infected with hepatopancreatic parvovirus (HPV) and monodon baculovirus (MBV). Again, no new data related to production losses were presented. That left only 3 references and one anecdotal statement dealing with production issues that would be relevant to OIE listing criterion 4b from 2010 onward.

One of these 3 remaining references was a review published by D.V. Lightner in 2011, but that paper does not refer to negative production impacts due to IHHNV from 2010 or 2011, so it is irrelevant to the analysis. Thus, only 2 publications remained in the OIE document that proposed evidence of losses to IHHNV between 2010 and 2020. There was also one anecdotal statement (unsupported by documentation) regarding losses to IHHNV at an indoor shrimp production facility in the United Kingdom. An analysis of these 3 issues follows.

The two publications cited were Jagadeesan et al. 2019 (Jagadeesan et al., 2019) and Sellars et al. 2019 (Sellars et al., 2019). Their conclusions are based mainly on positive PCR results for IHHNV and do not specify commercial production farm losses. The paper by Jagadeesan et al. contained results from a survey of 350 farms cultivating the whiteleg shrimp *P. vannamei* in India. From 30/350 farms that gave positive PCR test results for IHHNV, 14/30 were “IHHNV positive farms with no deformities, decreased growth and size variation”. In addition, the description of results from laboratory challenge by feeding with IHHNV positive tissue was, “The animals exhibited normal feeding, swimming behaviour. No specific clinical signs pertaining to this disease were observed in the animals.” In addition, the histological analysis did not show Cowdry Type A inclusions (i.e., eosinophilic intranuclear inclusions separated from an unstained space by marginated chromatin) characteristic of IHHNV. Also, *in situ* hybridization results were not accompanied by photomicrographs of adjacent tissue sections stained with hematolylin and eosin (H&E) or by photomicrographs of positive and negative controls. Finally, no data were presented regarding comparative production results from the studied farms. Given that approximately half of the 30 (14/30) ponds were IHHNV-positive but showed no gross signs of disease while the other 16/30 were positive with signs of disease would suggest that IHHNV was not statistically associated with the disease signs observed. Finally, no data were provided regarding analysis for other possible causes of deformation including abdominal segment deformity disease (ASDD) (Sakaew et al., 2008) or causes of retarded growth such as Laem Singh virus (LSNV) (Sritunyalucksana et al., 2006) and hepatopanceatic microsporidiosis (HPM) caused by *Enterocytozoon hepatopenaei* (Tourtip et al., 2009). All in all, this publication does not provide relevant information to support Criterion 4b. Finally, the study covered the period 2013-2018 (6 years), during which period we could not find other reports concerning RDS or other serious losses to IHHNV in production ponds in India. Thus, this publication alone, lacking substantive information on significant production losses, cannot be used to support the contention that IHHNV resulted “in significant consequences e.g., production losses, morbidity or mortality at a zone or country level”.

The publication by Sellars et al. (2019) was a report from research ponds cultivating *P. monodon* rather than commercial farm production ponds, and it contained no information related to the impact of IHHNV on farmed shrimp in Australia. In addition, *P. vannamei* is not cultivated in Australia and, as far as we know, there have been no reports of significant production losses associated with IHHNV in the *P*. *monodon* cultivated there. This is particularly relevant, because wild, captured broodstock are used to produce the postlarvae to stock commercial farms and there is no indication that the broodstock are even tested for IHHNV before use. This is despite the fact that IHHNV is known to be endemic and occurs at approximately 30% in Australian *P. monodon* (Huerlimann et al., 2018). If it were responsible for significant production farm losses, there should be some reports since at least 2004.

In the study by Sellars et al. (2019), the post larvae used to stock the experimental ponds were tested for IHHNV and two other viruses (yellow head viruses Type-2 and Type-7 known to occur naturally in Australian shrimp) by PCR only. However, no supporting histological analysis was carried out to confirm the presence or severity of IHHNV lesions. In addition, no PCR tests or histological analyses were carried out for monodon baculovirus (MBV) [now called Penaeus monodon nudivirus or PmNV (Yang et al. 2014)] or hepatopancreatic parvovirus (HPV) [decapod hepanhamaparvovirus or DHPV) (Srisala et al., 2021). Both are well known to be causes of retarded growth in *P. monodon* (Flegel, 2012; Thitamadee et al., 2016) and to sometimes accompany IHHNV in dual or triple infections (Flegel et al., 2004). In addition, the “losses to IHHNV” in the report from 1600 m^2^ experimental ponds the results were extrapolated to 1 hectare ponds (10,000 m^2^). This gave projected production at 155 days of culture as 9.0 and 10.1 t/ha (mean shrimp weight approximately 27 g) from the ponds with high IHHNV loads and 12.3 and 14.2 t/ha (mean shrimp weight approximately 33 g) in ponds with low IHHNV loads. These results contrast sharply with the lower, 6.0-7.5 t/ha normally expected at around 140 days with mean shrimp weights of approximately 30-35 g (Marín‐Riffo et al., 2021; West et al., 2019). In Thailand, excellent harvests of *P. monodon* consist of mean size 30 g at around 8 t/ha in approximately 120 days.

Thus, the projected 1 ha yield even for the high-level IHHNV experimental ponds would have been considered much better than normal based on usual farming results (Marín‐Riffo et al., 2021), while that projected for the low-level IHHNV ponds (12.3 and 14.2 t/ha) has rarely if ever been reported from anywhere as far as we know for ponds that are not progressively harvested. In addition, cultivation also lasted longer (155-169 days) to reach mean shrimp weights of 30-35 g usually reached in normal Australian ponds around 140 days. Furthermore, the mean weights at 86 days for the shrimp in the two pond groups, “mean weights (7.27 ± 4.06 to 7.83 ± 4.01) were similar across all 4 ponds” and were not significantly different. In addition, the SD values gave coefficients of variation (CV) for weight of 56% and 51% for the IHHNV high-load and low-load ponds, respectively. These are abnormally high CV values indicative of abnormally high size variation in both ponds as opposed to expected CV’s below 30% for normal ponds, but we do not know the size distribution curves for the ponds, so we cannot compare whether the curves for the two pond types were shifted towards non-profitable small sizes.

Altogether, the results from these experimental ponds do not relate to the normal experience, even for the low-IHHNV-level experimental ponds. Thus, they should not be used to support the contention that IHHNV has a significant negative impact on farmed shrimp production. To support this contention, it is important that there have been no reports from Australia regarding significant farmed shrimp production losses caused by IHHNV since at least 2004, despite knowledge that the virus is prevalent in captured wild shrimp (Huerlimann et al., 2018) used to produce post larvae for stocking Australian shrimp farms. Thus, the publication by Sellars et al. (2019 cannot be used to support the contention that IHHNV has resulted “in significant consequences e.g., production losses, morbidity or mortality at a zone or country level”.

The OIE review did not include similar reports on the occurrence IHHNV in exotic *P. vannamei* cultivated in Korea and proposed to have arisen from imported shrimp from retail outlets (Kim et al., 2011; Kim et al., 2012; Kim et al., 2020; Park et al., 2020). First of all, these PCR-based studies (like those above) presented no histopathological evidence to confirm IHHNV pathology or absence of pathology from other known or unknown pathogens. Since Korea does not have native *P. vannamei* the ponds must have been stocked with imported PL or PL derived from *P. vannamei* cultivated in Korea. However, no evidence was presented to prove that the broodstock used to produce the PL or the PL used to stock the shrimp ponds were free or IHHNV or other pathogens when stocked in the ponds. These reports cannot be used as evidence to support listing of IHHNV.

With respect to the OIE report reference to anecdotal evidence of IHHNV “disease outbreaks” at indoor shrimp production facilities for *P. vannamei* in the UK, we could find no published study available to assess the event. Given the uncertainty regarding this “anecdotal” information, it is inappropriate that it be included as major evidence in a discussion related to IHHNV impact on worldwide shrimp production. On the other hand, we have received unconfirmed information that the outbreaks arose due to failure to detect IHHNV in the imported *P. vannamei* shrimp stock. Since the stock was apparently destroyed under supervision of the UK competent authority, we hope that the owner was compensated for the loss.

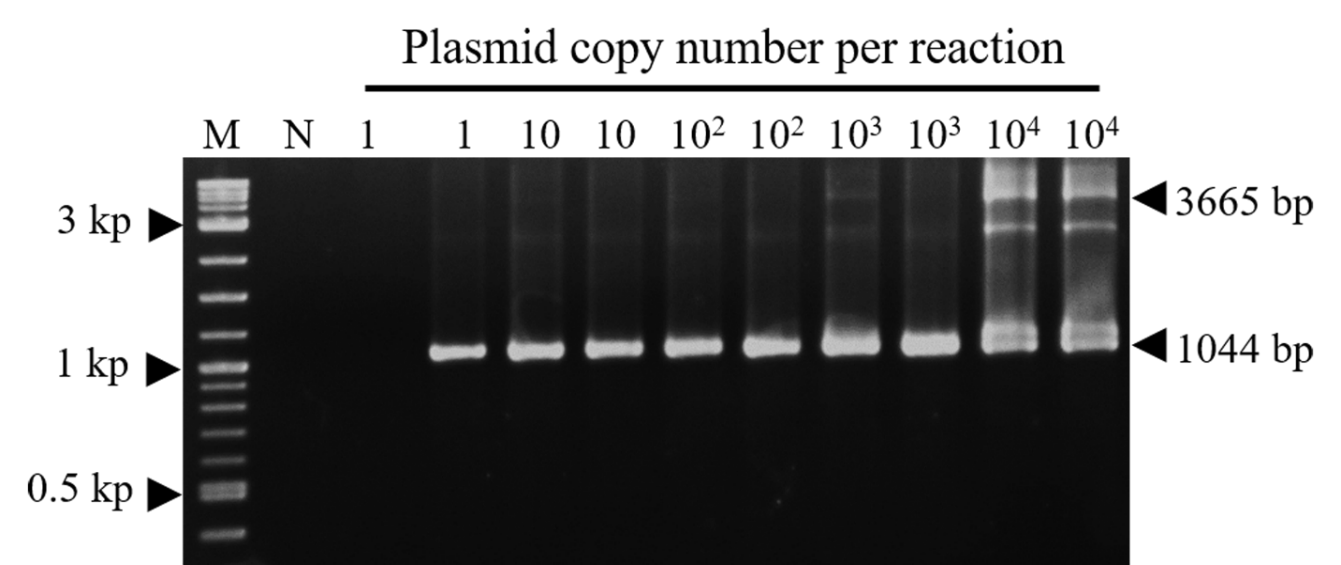

**Supplementary Figure S1.** Evaluation of the sensitivity of long-amp nested PCR. The long-amp nested PCR was tested by using plasmid DNA containing IHHNV sequence as a template. The sensitivity of detection was reached to 10 copies IHNNV genome based on nested PCR detection.

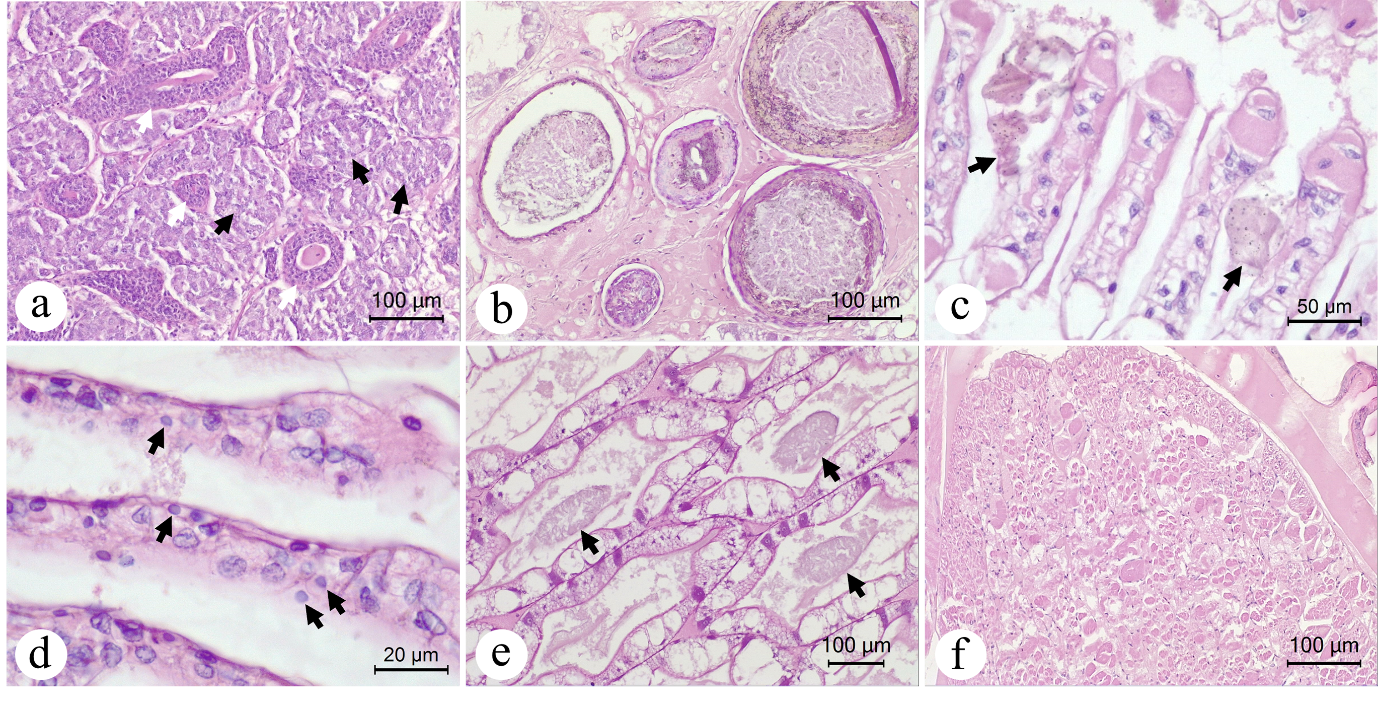

**Supplementary Figure S2.** Examples photomicrographs of the histopathologies for Pond Set 1. (a) Lymphoid organ spheroids (block arrow) and normal LO tubule (white arrow). (b) Bacterial granulomas in HP. (c) Gill fouling. (d) Gill pyknotic nuclei. (e) ATM. (f) Muscle lesions

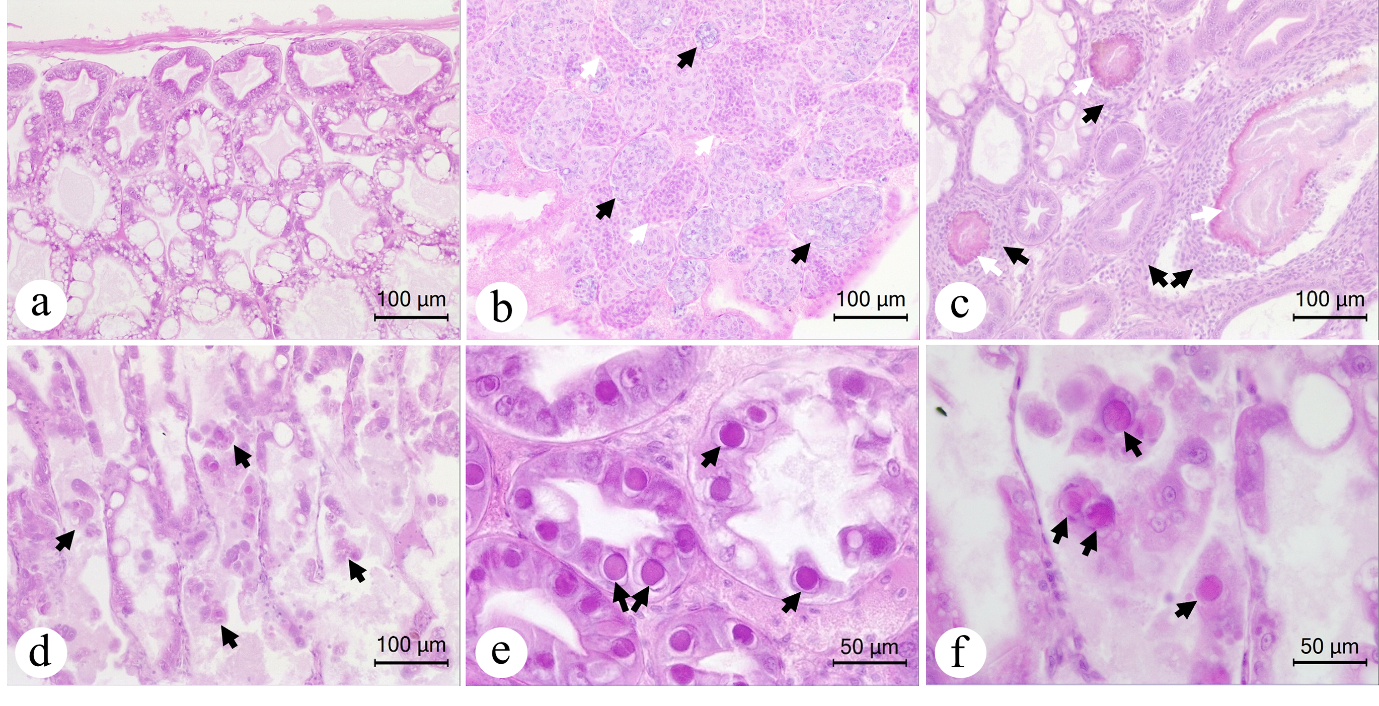

**Supplementary Figure S3**. Examples photomicrographs of the histopathologies for Pond Set 2. (a) Normal HP. (b) Lymphoid organ, spheroids (block arrow) LO tubule (white arrow). (c) HP bacterial lesions with granuloma (white arrow) and hemocyte aggregation (block arrow). (d) HP sloughing cells. (e) HPV inclusion in HP. (f) MBV inclusion in HP.
